# Constitutive activity of an atypical chemokine receptor revealed by inverse agonistic nanobodies

**DOI:** 10.1101/2024.11.04.621790

**Authors:** Claudia V. Perez Almeria, Omolade Otun, Roman Schlimgen, Thomas D. Lamme, Caitrin Crudden, Noureldine Youssef, Lejla Musli, Shawn Jenjak, Vladimir Bobkov, Julia Drube, Carsten Hoffmann, Brian F. Volkman, Sébastien Granier, Cherine Bechara, Marco Siderius, Raimond Heukers, Christopher T. Schafer, Martine J. Smit

## Abstract

Chemokine stimulation of atypical chemokine receptor 3 (ACKR3) does not activate G proteins but recruits arrestins. It is a chemokine scavenger that indirectly influences responses by restricting the availability of CXCL12, an agonist shared with the canonical receptor CXCR4. ACKR3 is upregulated in numerous disorders. Due to limited insights in chemokine-activated ACKR3 signaling, it is unclear how ACKR3 contributes to pathological phenotypes. One explanation may be that high constitutive activity of ACKR3 drives non-canonical signaling through a basal receptor state. Here we characterize the constitutive action of ACKR3 using novel inverse agonistic nanobodies to suppress basal activity. These new tools promote an inactive receptor conformation which decreased arrestin engagement and inhibited constitutive internalization. Basal, non-chemotactic, breast cancer cell motility was also suppressed, suggesting a role for ACKR3 in this process. The basal receptor activity in pathophysiology may provide a new therapeutic approach for targeting ACKR3.

## Introduction

Atypical Chemokine Receptor 3 (ACKR3, formerly CXCR7) is a β-arrestin-biased chemokine receptor^1^ that lacks detectable G protein activation in most cell types (with the exception of primary rodent astrocytes and human glioma cells)^1–3^. Activation of the receptor leads to phosphorylation of C-terminal serine and threonine residues by GPCR kinases (GRKs)^4–7^. These modifications are critical for coordinating arrestin coupling. ACKR3 is best described as a scavenger, where its primary function is to regulate the extracellular concentrations of ligands and restrict the ligand availability for canonical receptor activation. The receptor shares chemokine ligands with both CXCR4 (CXCL12) and CXCR3 (CXCL11), both of which drive cell migration along chemokine gradients. Scavenging by ACKR3 therefore indirectly supports chemotaxis by generating directional information and preventing overstimulation and desensitization of CXCR4 or CXCR3. This regulatory activity is dependent on GRK phosphorylation, but not arrestin engagement^4,6^, suggesting the receptor might better be regarded as a GRK-biased receptor^5^. Besides chemokines, ACKR3 is activated by opioid peptides (BAM22, enkephalins, and dynorphins)^8,9^ and pro-adrenomedullin derivatives (adrenomedullin and PAMP-12)^10,11^. The wide range of natural ligands binding ACKR3 suggests a flexible binding pocket and a promiscuous receptor^12^. ACKR3 is involved in many physiological functions^13^ as well as in a plethora of pathophysiological processes, including inflammatory^14^, autoimmune^15^, and neurodegenerative diseases^16^ in addition to different types of cancer^17^. ACKR3 overexpression is associated with neurodegeneration in the central nervous system and poor cancer prognosis, while it provides a cardioprotective role in cardiovascular diseases^18^.

GPCR signaling plays an important role in controlling various cancer hallmarks^19^. The CXCL12-CXCR4-ACKR3 axis contributors are key to cancer cell migration, survival, and proliferation^20,21^. Enhanced ACKR3 expression in numerous cancer types (e.g. glioma, lung, breast, colorectal, lymphoma), has been associated with the shaping of CXCL12 gradients, by internalizing with the chemokine and recycling the receptor back to the plasma membrane^22^. Through this mechanism, ACKR3 appears pivotal for tumorigenesis, angiogenesis, cell adhesion, and tumor growth^23–27^. Despite ACKR3’s evident role in cancer development, the specific downstream signaling pathways modulated by this receptor are still unclear. Numerous studies have suggested that CXCL12-stimulated ACKR3 signals via β-arrestin-dependent pathways activating ERK and AKT^1,28–30^. However, recent reports indicate that these may be ascribed to background CXCR4 signaling by G proteins^5,31^.

In addition to chemokine-induced responses, ACKR3 displays considerable constitutive activity in the apo (empty) receptor state. The receptor readily interacts with arrestin in the absence of stimulation, both in cells^12^ and detergent purified form^32^. Without a ligand bound, ACKR3 flexibly interconverts between active and inactive conformation^33^, which leads to basal phosphorylation by GRKs that coordinates arrestin binding^4^. Additionally, the receptor constitutively internalizes by mechanisms independent of C-terminal phosphorylation^4,5^. This internalization contributes to scavenging, but is unable to dynamically respond to large fluxes in chemokine concentration^34^. This high level of constitutive activity may explain difficulties in antagonizing the receptor, as only a handful of inhibitors have been described^32,33,35,36^. It is unknown whether the constitutive activity of ACKR3 contributes to other cellular processes and if these deviate from chemokine-induced responses. Different signaling states for constitutive and agonist-stimulated activation have been observed for a virally-encoded chemokine receptor, US28^37^. Uncharacterized signaling by the apo-receptor may play an unappreciated role in ACKR3 physiology and pathophysiology.

In order to resolve the constitutive mechanisms of ACKR3 function, we developed new nanobody-based inhibitors to suppress basal activation of the atypical receptor. Nanobodies, also known as single-domain antibodies or VHHs, are the variable domains from heavy chain-only antibodies found in the Camelidae family. Nanobodies display high affinity and specificity for their target and tend to interact with non-linear, 3-dimensional epitopes. These features make them ideal molecules for targeting and stabilizing GPCRs in specific conformational states^38–41^, which may be particularly important for a promiscuous protein like ACKR3. Using advanced structural dynamics methods, we showed that the nanobodies stabilize inactive receptor conformations that correlate with inhibited basal engagement with arrestins and constitutive internalization. Inhibition of receptor constitutive activity led to slower cell motility. These data highlight the potential consequences of ACKR3 basal activity.

## Results

### Basal ACKR3 engagement with arrestins is suppressed by inverse agonistic nanobodies

An antagonistic nanobody targeting ACKR3, VUN701, was recently characterized^42^. Here, we present two additional nanobodies, VUN700 and VUN702, which were not previously characterized pharmacologically. All three nanobodies bind the receptor extracellularly and compete with CXCL12 (Supplementary Fig. 1, Supplementary Table 1). Due to the bulky and relatively large binding interface of nanobodies and chemokines, nanobodies sterically prevent co-binding. As a consequence, nanobodies binding to extracellular domains of chemokine receptors generally act as antagonists^39,43–45^, though some are agonists^46^ or have been engineered to activate receptors^47^. To resolve the pharmacological effects of these molecules, ACKR3 engagement with arrestin was tracked by BRET between the receptor with a C-terminal nanoluciferase (ACKR3-Nluc) and β-arrestin2 C-terminally tagged with mVenus (β-arr2-mV), following addition of CXCL12 agonist or the nanobodies. Activation by the agonist CXCL12 led to a robust increase in BRET ratio, indicating a recruitment of arrestin to the receptor in HEK293T cells (Fig. 1A, B). When the cells are treated with neutral antagonist VUN701, no change in association of ACKR3 with β-arrestin was detected, consistent with its previous pharmacological classification^48^. Interestingly, VUN700 and VUN702 both decreased the BRET ratio between ACKR3 and β-arrestin2 below the measured basal interaction. This suggests that these nanobodies are acting as inverse agonists and suppressing the previously reported constitutive ACKR3 activity^12,33^. Similar results were observed with β-arrestin1 recruitment (Supplementary Fig. 2).

**Figure 1.**
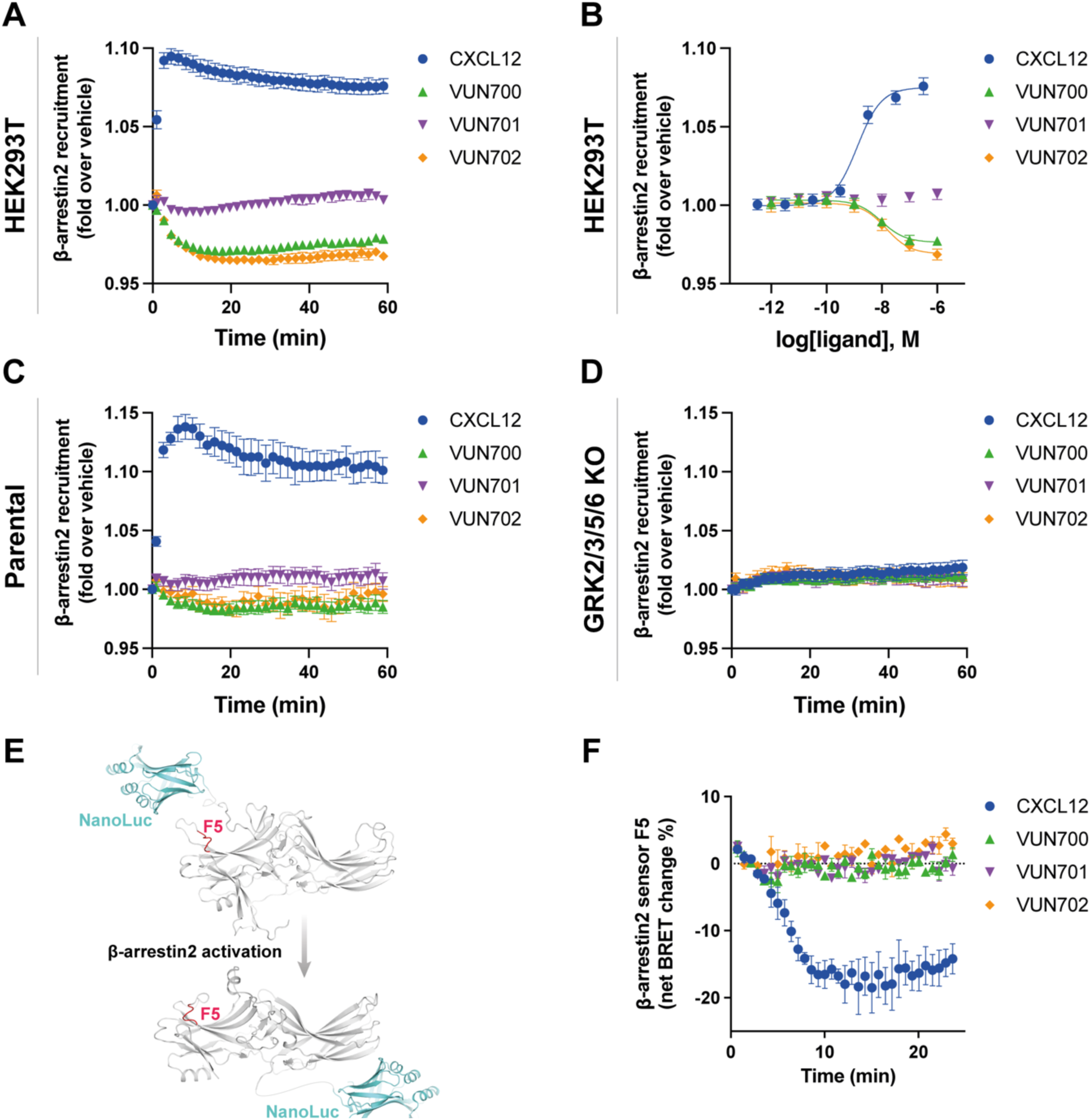
ACKR3 nanobodies suppress basal β-arrestin2 recruitment. **A-B)** Recruitment of β-arr2-mV to ACKR3-Nluc measured by BRET **(A)** Time-dependent change in BRET over 60 min with either 316 nM of CXCL12 (blue circle) or 1 μM of nanobody (VUN700 (green triangle), VUN701 (purple inverted triangle), VUN702 (yellow diamond)) and **(B)** dose response curves of CXCL12 or nanobodies at 60 min recorded at 37°C in HEK293T cells. **C-D)** β-arrestin2 recruitment measured by BRET in agonist mode between donor ACKR3-Nluc and β-arr2-mV in **(C)** parental HEK293 or in **(D)** GRK2/3/5/6 KO HEK293 cells. **E)** Schematic illustration of BRET-based FlAsH-tagged (CCPGCC) sensor F5 (between residues 156 and 157) on β-arrestin2. **F)** Time-resolved changes in the NanoBRET β-arrestin2 conformational biosensor F5 signal upon ACKR3 activation, following the addition of 316nM of CXCL12 or 1 μM of VUN700, VUN701 or VUN702 at 37°C in parental HEK293 cells. Data are shown as the average ± SD of three independent experiments performed in technical triplicates.

ACKR3 requires phosphorylation by GRKs to engage β-arrestins in response to CXCL12^5,6,49^, while constitutively active arrestin can interact with unmodified apo-ACKR3 in vitro^32^. To ascertain the role of GRK phosphorylation in basal association in cells, arrestin recruitment was also tested in CRISPR-knockout cells of the four ubiquitously expressed GRKs, GRK2, 3, 5, and 6 (GRK2/3/5/6 KO)^50^. In these cells, the response to CXCL12 was completely abolished, consistent with previous results (Fig. 1C, D). Additionally, the inverse agonistic effects of VUN700 and VUN702 disappeared in the absence of GRKs. Together, these data suggest that the basal arrestin association to ACKR3 is GRK-dependent and likely reflects the phosphorylation of constitutively active receptors by these kinases.

To further resolve differences between agonist-induced and basal arrestin engagement with ACKR3, the conformational changes within the arrestins were monitored using Nluc/FlAsH arrestin intramolecular BRET biosensors^51,52^. These sensors report subtle differences in arrestin conformations, corresponding to the active conformations arrestin adopts due to its interaction with GPCRs (Fig. 1E). As reported by the FlAsH 5 (F5 sensor), activation by CXCL12 promoted a robust decrease in BRET, indicating the adoption of an active arrestin state. None of the nanobodies produced a change in the signal from this sensor. Similar responses were observed for two other arrestin conformational sensors with different FlAsH positions (Supplementary Fig. 3). This implies that either the basal interaction of ACKR3 and β-arrestin2 does not induce a conformational change in arrestin or the reversion is not detectable by these sensors.

### Distinct ACKR3 conformational states are stabilized by antagonist and inverse agonist nanobodies

The inhibition of basal arrestin interactions of ACKR3 by VUN700 and VUN702 suggests that these nanobodies act as inverse agonists, possibly by inducing a more inactive-like receptor conformation. Two structural dynamics methods were used to determine how the nanobodies were specifically altering the conformation of ACKR3; nuclear magnetic resonance (NMR) and hydrogen-deuterium exchange mass spectrometry (HDX-MS).

First, ^13^CH_3_-e-Met labelled ACKR3 was purified and analyzed by NMR spectroscopy^53^. As previously described^48^, two of ACKR3’s eight native methionines (M212^5×39^ and M138^3×46^, GPCRdb nomenclature in superscript^54^ Supplementary Fig. 4) can be used to track receptor conformational dynamics at the ligand binding site and in the intracellular region, respectively (Fig. 2A). At first glance, the NMR analysis of ACKR3 bound to VUN700 or VUN702 results in similar spectra to that of ACKR3 bound to the neutral antagonist VUN701 (Fig, 2B). However, overlaying the spectra from the different ACKR3-nanobody complexes does reveal subtle shifts in the peaks for M212^5×39^ (Fig. 2C) and M138^3×46^ (Fig. 2D). Upon CXCL12 binding, the M212^5×39^ position was previously shown to be in a gauche rotameric state with a peak at 16.25 ppm. This shifted downfield to 17.0 ppm with the neutral antagonist VUN701^48^. Binding of VUN700 or VUN702 further altered the M212^5×39^ peak as compared to the ACKR3-VUN701 complex (Fig. 2C). In all three complexes, the ^13^C position of ∼17.0 ppm is consistent with rotamer averaging and the absence of stabilizing interactions at the M212^5×39^ position. In contrast, the M138^3×46^ position showed a slight downfield shift in the ^13^C and ^1^H dimensions for VUN701, compared to the inverse agonists (Fig. 2D). Given the previous evidence that M138^3×46^ exists as a mixture of active and inactive states, the shift along this line suggests that VUN700 and VUN702 binding shift the ACKR3 conformational equilibrium relative to VUN701, potentially indicating a more “OFF” state of the receptor (Figure 2D).

**Figure 2.**
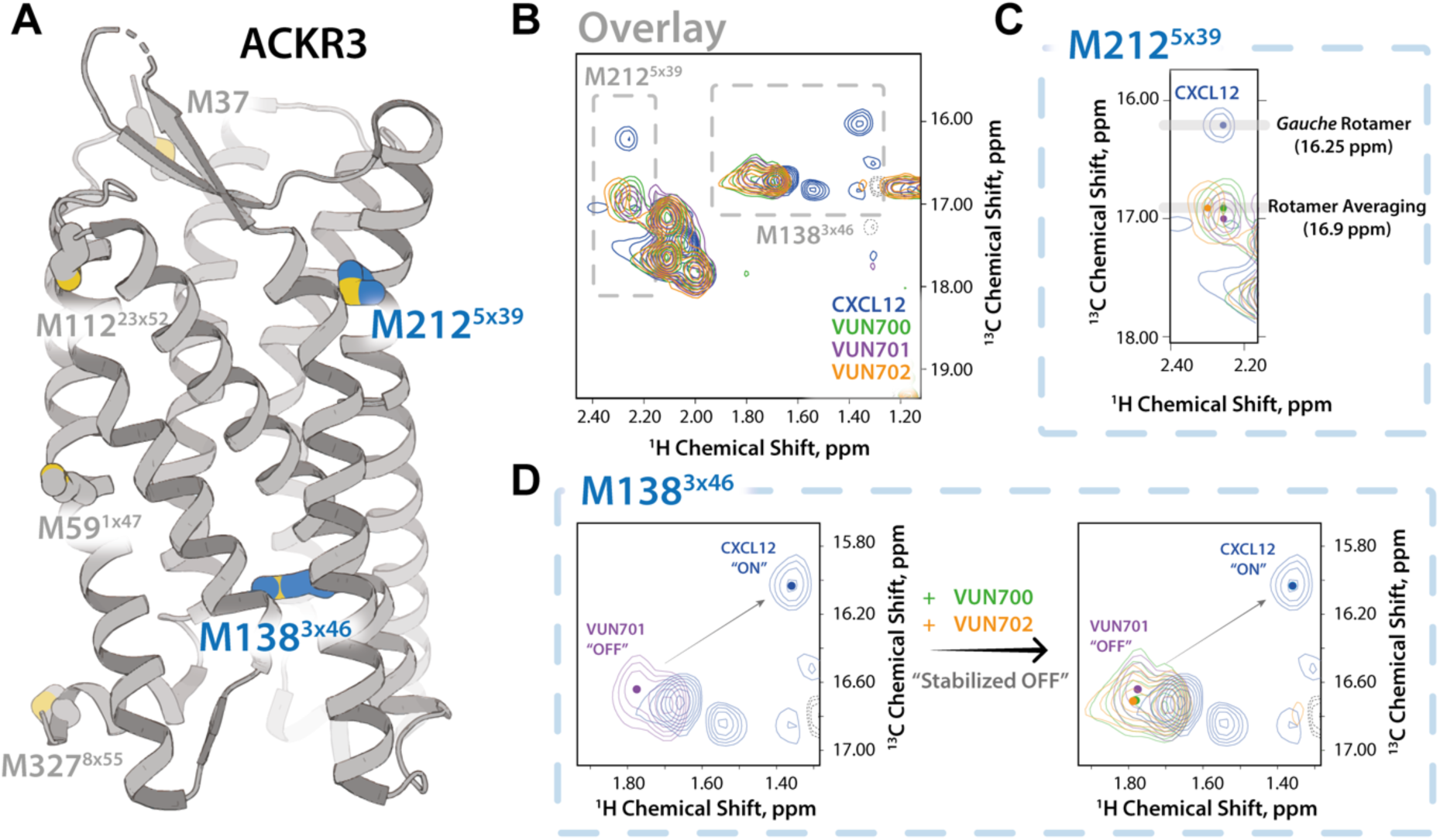
NMR-based structural characterization of ACKR3 upon nanobodies VUN700 binding reveals a relatively more pronounced “OFF” state of ACKR3 than VUN701-bound state. **A)** ACKR3 structure (7SK6 PDB)^12^ with NMR peaks M138^3×46^ and M212^5×39^ depicted. **B)** Overlay of M212^5×39^ peaks from all nanobodies and CXCL12 ligand-bound ACKR3 complexes. **C)** Overlay of M138^3×46^ peaks in all nanobodies and CXCL12 ligand-bound states. Upfield peak positions (^1^H: ∼1.3 ppm) of M138^3×46^ among agonist-bound states supports ring-current shifts due to aromatic side chain interactions.

To further structurally substantiate the subtle conformational changes observed by NMR, induced by the inverse agonist and antagonist nanobodies, HDX-MS was performed to track changes in the rate of isotopic exchange between amide hydrogens on ACKR3 and deuterium in the solvent. The exchange rate depends on solvent accessibility and hydrogen bonding networks, and comparing these rates provides insights into changes to the protein conformational state and protein-binding interfaces. This technique has recently been optimized for ACKR3 to monitor conformational changes due to small molecule ligands^32^.

Using differential HDX (ΔHDX) analyses, we compared the unbound (apo) and nanobody-bound states of ACKR3 (Fig. 3A, Supplementary Fig. 5A). Binding of the nanobodies protected the extracellular face from deuteration, confirming the nanobody binding interface proposed from CXCL12 competition assays (Supplementary Fig. 1). Differences in deuteration were localized to peptides corresponding to the orthosteric (CXCL12) binding pocket at the N-terminus (N-term, residues 27-33) and TM5 (residues 204-211) which displayed large protection (respective ΔHDX of up to 15% and 30% for each nanobody) (Supplementary Fig. 5B). The nanobodies significantly protected peptides in the N-terminus and extracellular loops (ECLs) of ACKR3 that correspond to the CXCL12 binding interface^12^, suggesting that the nanobodies bind similarly to these receptor regions in agreement with our previous model^42^.

**Figure 3.**
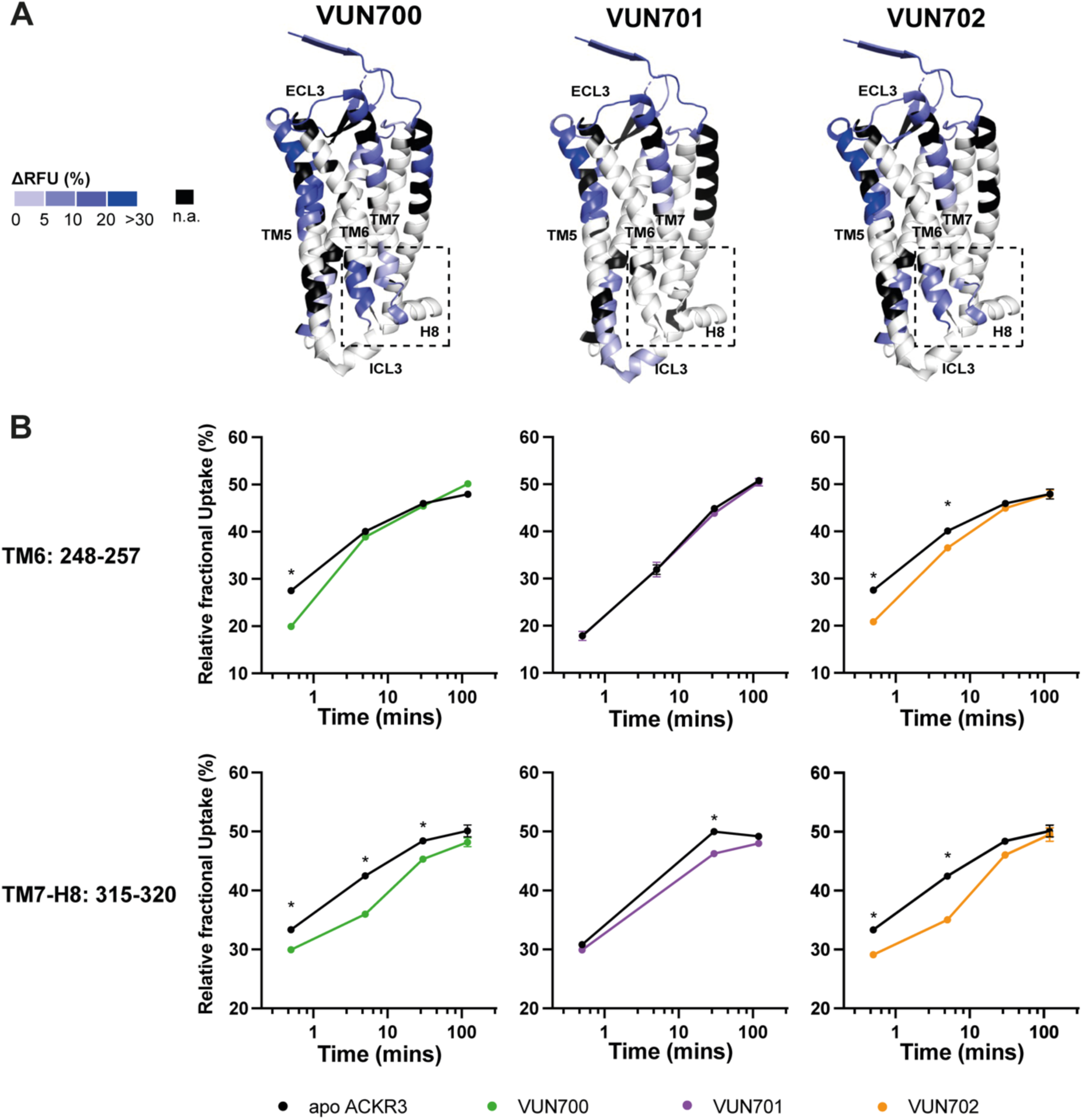
Conformational changes in ACKR3 induced by nanobody binding. **A).** Structural representation of the % differential relative fractional uptake (ΔRFU) data (apo ACKR3 – Nb-bound ACKR3) mapped onto the cryo-EM structure of ACKR3 (PDB:7SK5). This depicts reproducible and statistically significant ΔHDX over 120 minutes deuteration. The degree of ΔHDX (% ΔRFU) ΔRFU is represented according to the color scale. Black regions represent those with no sequence coverage. **B)** Deuterium uptake plots showing time-dependent change in RFU for ACKR3 peptides on the intracellular side upon nanobodies binding, compared to apo ACKR3 (in black). Uptake represents the average and SD of three technical replicates from one biological preparation of ACKR3. Data is representative of three biological replicates. Statistically significant changes were determined using Deuteros 2.0 software^90^ (*, p ≤ 0.01).

While the overlapping interacting sites between CXCL12 and the nanobodies with ACKR3 explain their competitive binding mode, nanobody binding also induced conformational changes to the intracellular side of the receptor (Fig. 3B). It is known that the position of the cytoplasmic ends of TM6 and TM7 reflect the active state of GPCRs including ACKR3^12,32,33^. Only slight differences were observed for these regions with the neutral antagonist VUN701 bound. In contrast, both inverse agonists VUN700 and VUN702 showed robust protection at the intracellular face of TM6, residues 248^6×31^ to 257^6×40^. Likewise, the inverse agonists induced protection at the linker region connecting TM7 to H8, residues 315^7×53^ to 320^8×48^, whilst VUN701 did not. These differences in HDX protection suggest that the inverse agonism observed for VUN700 and VUN702 is due to the promotion of an inactive ACKR3 conformation, while the neutral antagonist VUN701 does not impact the basal state of the receptor. Both results are consistent with the biological responses observed in Fig. 1. Together with the NMR-based analysis, this HDX analysis suggests a unique conformational state for the inverse agonist-bound ACKR3 compared to when bound to an antagonist.

### Inverse agonistic nanobodies trap ACKR3 at the plasma membrane

Constitutive internalization of ACKR3 contributes to chemokine scavenging^4,7,22,55^ and is independent of receptor phosphorylation^5^. Therefore, we examined whether the inverse agonism displayed by these nanobodies had functional consequences on receptor trafficking from the plasma membrane (PM) to early endosomes. First, we examined how the different nanobodies modulate ACKR3 internalization by monitoring the presence of ACKR3 at the plasma membrane with flow cytometry (Fig. 4A). CXCL12 internalized 25% of ACKR3 after 15 min exposure of cells, and after 45 min ACKR3 returned back to basal surface levels. In contrast, all nanobodies induced an increased level of ACKR3 on the membrane over time. The inverse agonists VUN700 and VUN702 increased the receptor level on the membrane by up to ∼70% after 60 min of incubation. Interestingly, the neutral antagonist VUN701 also increased membrane presence of ACKR3, but to a lesser extent (∼50%) and with a delay (Fig 4A). We then investigated the subcellular trafficking of ACKR3 upon nanobody binding by employing BRET between ACKR3 and two different localization markers, mVenus-CAAX (mV-CAAX) for the plasma membrane and Rab5a-mVenus (Rab5a-mV) for the early endosomes^56^ (Fig. 4B, C). Upon CXCL12 binding, ACKR3 rapidly internalized away from the plasma membrane (Fig. 4B) and appeared in early endosomes (Fig. 4C). All three nanobodies inhibited constitutive internalization, causing the receptor to be retained at the membrane, consistent with the flow cytometry results. This was more prominent for the inverse agonists VUN700 and VUN702 than for neutral antagonist VUN701 (Fig. 4B). Similarly, VUN700 and VUN702 impaired the basal trafficking of ACKR3 to the early endosomes, while VUN701 had little effect (Fig. 4C). Taken together, nanobody binding blocks receptor intracellular trafficking by retaining the receptor on the membrane and this is more efficient for the inverse agonists than for the neutral antagonist VUN701.

**Figure 4.**
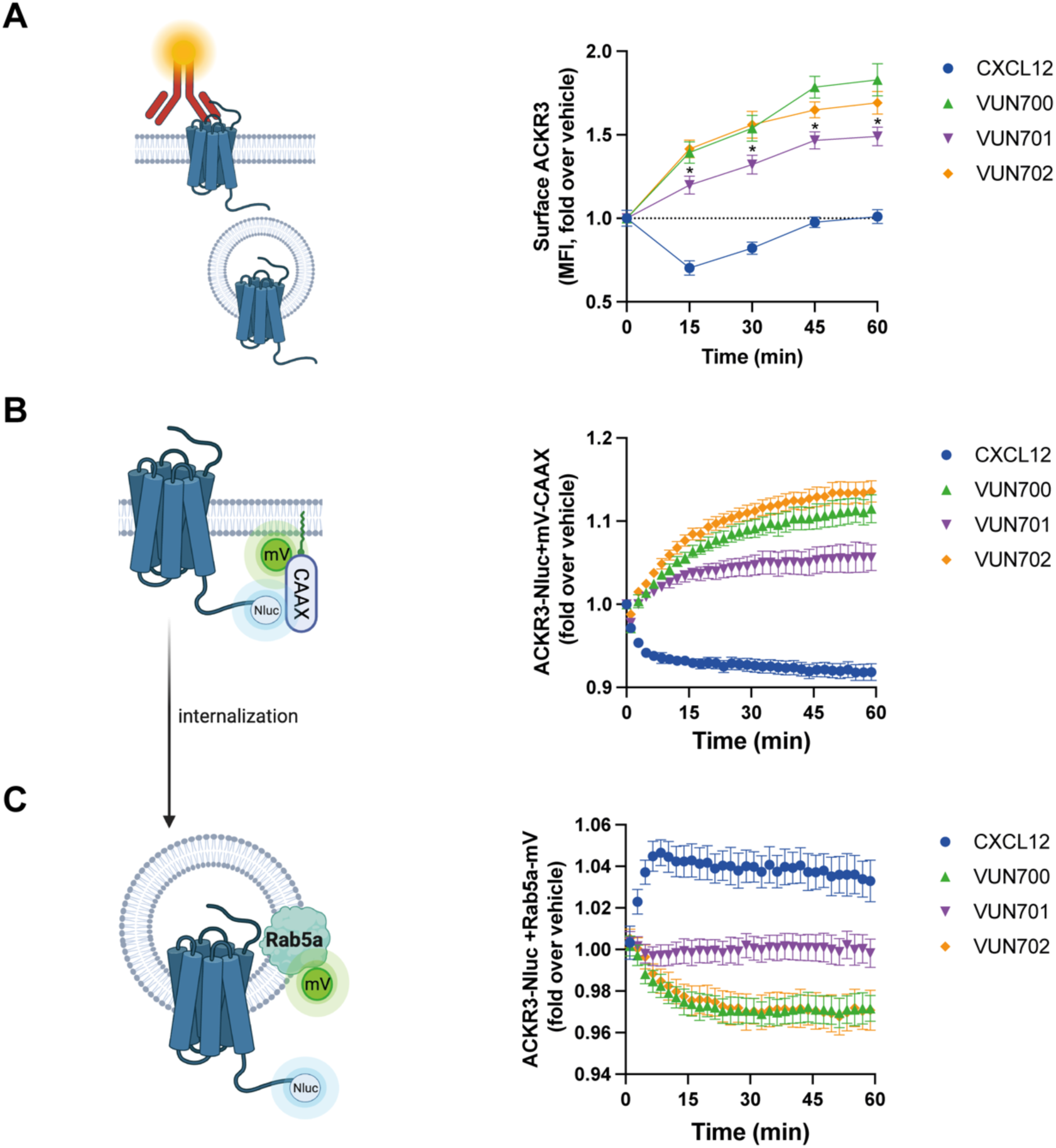
ACKR3 nanobodies differentially change the localization of ACKR3 by capturing ACKR3 at the membrane. **A)** Surface ACKR3 detected by flow cytometry upon 316 nM of VUN700 (green triangle), VUN701 (purple inverted triangle) or VUN702 (yellow diamond), or 100 nM CXCL12 (blue circle) over 60 min at 37°C in HEK293 cells. **B)** Time-dependent internalization measured by BRET, between donor ACKR3-Nluc with mV-CAAX over 60 min with either 316nM of CXCL12 or 1μM of VUN700, VUN701 or VUN702 at 37°C in HEK293T cells. **C)** Time-dependent ACKR3 localized in the early endosomes measured by BRET, between donor ACKR3-Nluc with Rab5a-mV, over 60 min with either 316nM of CXCL12 or 1μM of VUN700, VUN701 or VUN702 at 37°C in HEK293T cells. Data is shown as the average ± SD of at least three independent experiments performed in duplicates or triplicates. One-way ANOVA, multiple comparisons Dunnett test (* p<0.05).

### All nanobodies inhibit GRK-independent internalization of ACKR3

CXCL12-mediated internalization of ACKR3 is β-arrestin-independent^6^ but GRK-dependent^5^, while constitutive internalization is independent of both effectors. To determine if the nanobodies also impacted phosphorylation-independent internalization, the membrane presence of ACKR3 was observed by BRET in GRK2/3/5/6 KO HEK293 cells. In the parental cells, containing all GRKs, CXCL12 and the nanobodies showed the same order of effectiveness as shown in the HEK293T cells (Fig. 4B, 5A), with the inverse agonists VUN700 and VUN702 inducing greater retention on the membrane than the antagonist VUN701 (Fig. 5A, B). In the absence of GRKs, the internalization response from CXCL12 treatment is abolished, consistent with previous reports^5,49^. Unexpectedly, the effects of the inverse agonist and antagonist nanobodies were nearly identical without GRKs (Fig. 5C, D). The plasma membrane trapping effect by the nanobodies was independent of β-arrestins (Supplementary Fig. 6). These results suggest that constitutive internalization by ACKR3 can be divided into a phosphorylation-dependent component, which is suppressed by inverse agonism, and a phosphorylation-independent mechanism that is inhibited by both types of nanobodies tested here.

**Figure 5.**
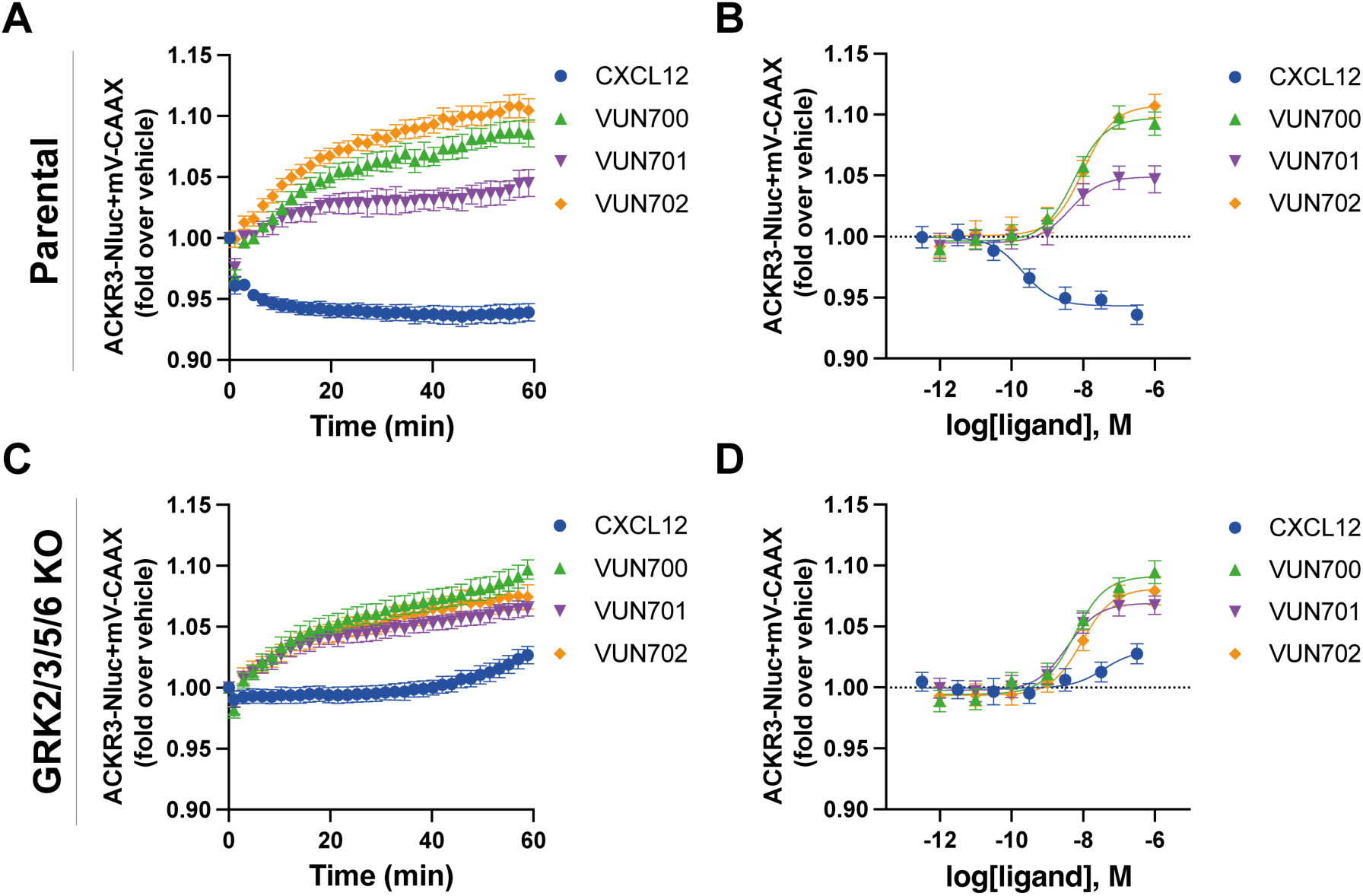
ACKR3 nanobodies mediate GRK-independent and dependent constitutive internalization. **A-D)** Internalization of ACKR3-Nluc measured by BRET to the PM probe mV-CAAX in (A-B) parental HEK293 or in (C-D) GRK2/3/5/6 KO HEK293 cells at 37°C. **(A, C)** Time-dependent change in BRET over 60 min with either 316 nM of CXCL12 (blue circle) or 1 μM of VUN700 (green triangle), VUN701 (purple inverted triangle) or VUN702 (yellow diamond) and **(B, D)** dose response curves of CXCL12 or nanobodies at 60 min. Data is shown as the average ± SD of four independent experiments performed in triplicate.

### Basal motility of metastatic breast cancer cells is reduced by ACKR3-directed inverse agonist nanobody

ACKR3 is reported to contribute to cancer cell migration^57^, but not due to activation by CXCL12^21,58^. Instead, we hypothesize that ACKR3’s constitutive activity might play a role in cell migration. To resolve the influence of ACKR3 on non-chemokine driven migration, the basal or random movement of metastatic breast cancer cells, MDA-MB-231, was tracked by live-single cell microscopy. MDA-MB-231 cells express relatively high levels of both ACKR3 and CXCR4 endogenously, which suggests an invasive phenotype and is associated with aggressive behavior^59^. This allows for examination of potential roles for ACKR3 in a relevant cellular context and in the presence of CXCR4. The cells showed considerable motility even without chemotactic stimulation (Fig. 6A). This basal motility was reduced when treated with either VUN700 or VUN701 (Fig. 6B). The degree of motility attenuation was quantified by comparing both the accumulated distance (total distance travelled) and Euclidean distance (straight line distance from cell starting point to end point). Both metrics decreased on average by ∼30% with inverse agonist VUN700 treatment. In the case of the neutral antagonist VUN701, only the accumulated distance was significantly impaired, while the change in overall position was only slightly altered (Fig. 6C). VUN400, a CXCR4 targeting nanobody that inhibits CXCL12 binding and CXCL12-induced chemotaxis^39^, had no effect on basal cell motility. Overall, these results suggest that basal activity of ACKR3 may influence MDA-MB-231 cancer cells motility in the absence of chemokine stimulation.

**Figure 6.**
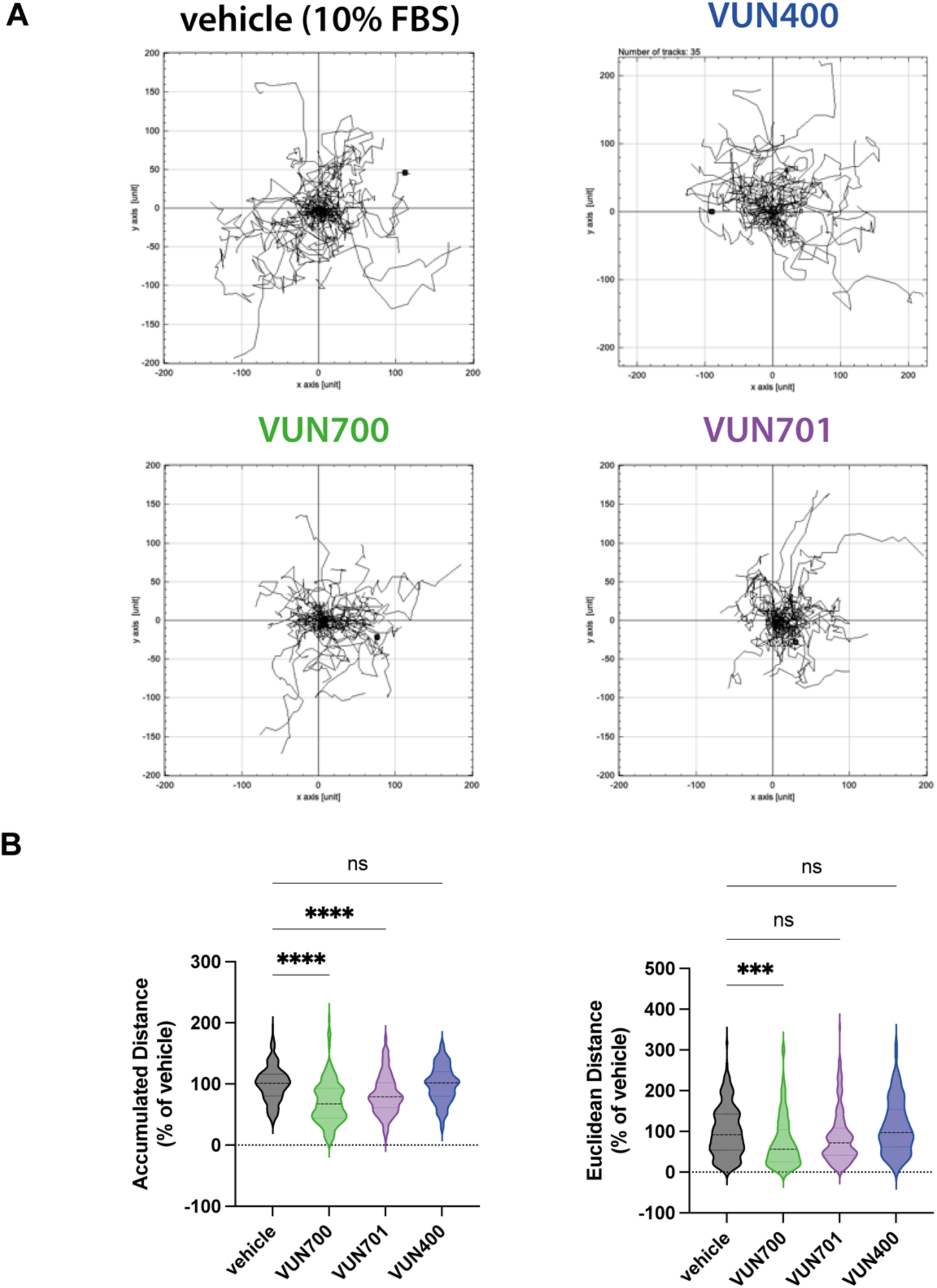
Basal motility in metastatic breast cancer cells is reduced upon inverse agonist nanobody targeting ACKR3. **A)** Representative basal motility paths of MDA-MB-231 metastatic breast cancer cells for 16h in 10% FBS, vehicle condition (in black) or with 100nM of VUN700 (in green), VUN701 (in purple) or VUN400 (in blue). **B)** Accumulated distance (total distance traveled) and Euclidean distance (Start to end point distance) for individual cells from each of the conditions in **A.** The positions of at least 120 cells, selected across three replicates (∼40 cells/repeat), were tracked and normalized to the vehicle condition. Significance was determined by one-way ANOVA Dunnett test (p<0.0005 (***), <0.0001 (****)).

## Discussion

ACKR3 is an atypical receptor that is best described as a chemokine scavenger. Although the receptor is implicated in many other physiological responses, they have not been explicitly tied to chemokine-mediated receptor activation or ligand scavenging. Here we present facets of ACKR3 constitutive activity with downstream responses using antagonistic and inverse agonistic ACKR3 nanobodies. The nanobodies had profound effects on the basal receptor events. While only the inverse agonistic nanobodies lead to a disruption of the arrestin-apo-ACKR3 complex, both inverse agonists and antagonists, albeit to a lower extent, suppressed constitutive internalization and trapped the receptor on the plasma membrane (Fig. 7). These effects appear to be due to subtle changes in the receptor’s conformational state and may manifest into attenuation of basal, or random, cellular migration. These data provide insight into hidden functions of the atypical receptor that are independent of chemokine receptor activation.

**Figure 7.**
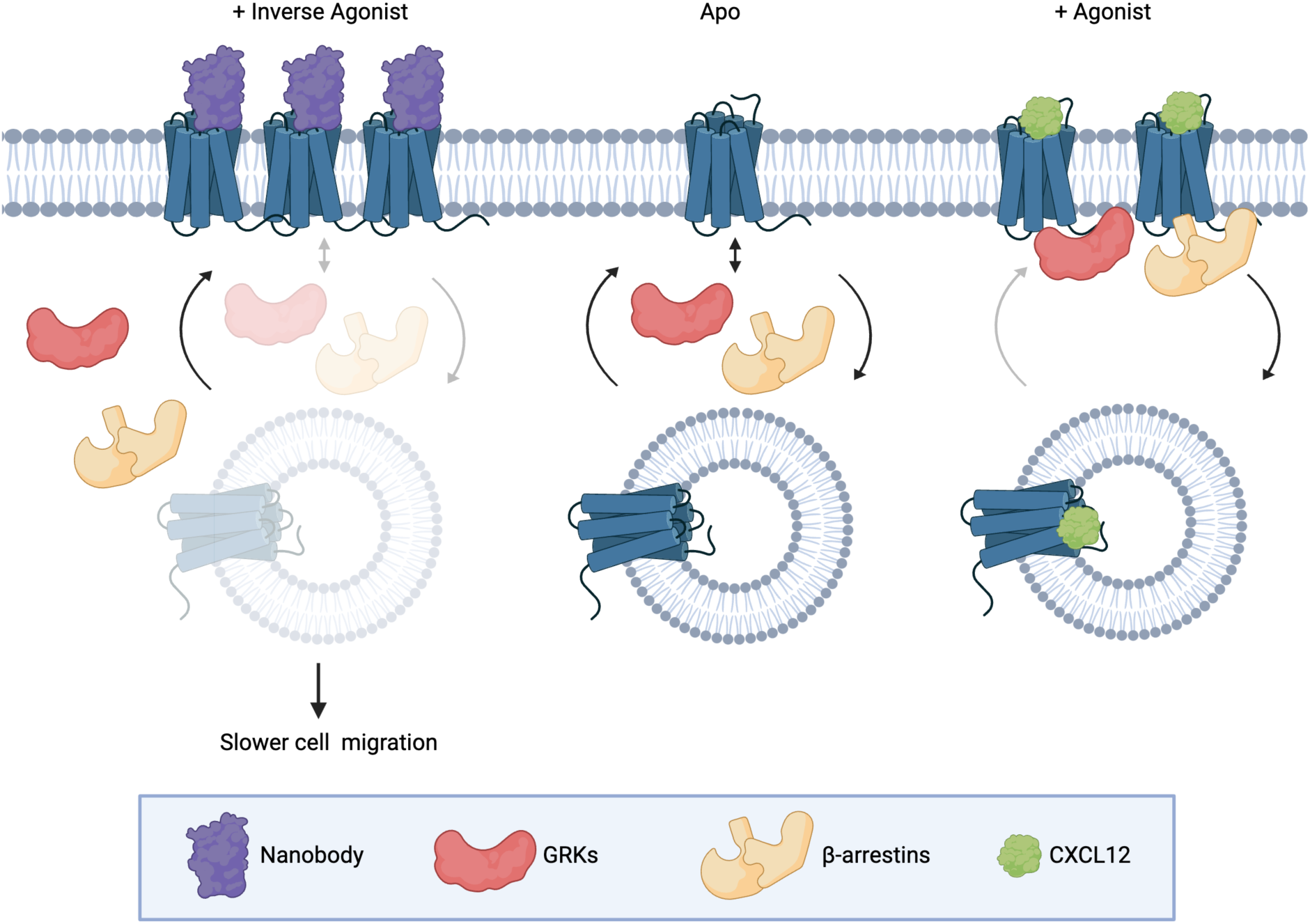
Constitutive activity of atypical chemokine receptor revealed by inverse agonistic nanobodies. Apo-ACKR3 is constitutively active, which leads to the receptor basal GRK phosphorylation, arrestin recruitment, and internalization. Stimulation with an agonist like CXCL12 fully activates ACKR3 and drives robust phosphorylation, arrestin complexing, and endocytosis. Inverse agonistic nanobodies suppress the constitutive activity of ACKR3, trapping the receptors at the cell surface and reducing interactions with arrestins or GRKs, thereby allowing GRKs and arrestins to be available for other GPCRs. These nanobodies also slow basal cell migration, suggesting a role for ACKR3 constitutive activity in cell motility.

Outward movement of TM6 is a common hallmark of GPCR activation^60,61^ and ACKR3 is no different^12,32,33^. The atypical receptor also displays extensive constitutive activity^12^ and readily adopts an active conformation in the absence of stimulation^33^. This constitutive activity drives basal GRK phosphorylation and subsequent arrestin engagement (Fig. 1). VUN700 and VUN702 act as inverse agonists to suppress basal β-arrestin engagement, while the previously characterized VUN701^48^ only blocked CXCL12-induced interactions. This suggests that different conformations are being stabilized by the nanobodies, leading to different effects on ACKR3 phosphorylation and arrestin interactions. Indeed, both HDX and NMR studies show slightly different conformations being promoted by the inverse agonists compared to VUN701. The protection observed at the cytoplasmic ends of TM6 and TM7 (Fig. 3B) as well as the chemical shifts of M138^3×46^ and M212^5×39^ (Fig. 2B, C) are consistent with inactive receptor states. The inverse agonists VUN700 and VUN702 appear to stabilize an even more inactive conformation than VUN701 (Fig. 2, 3). These structural observations are in line with the more profound effects observed for the inverse agonistic nanobodies on ACKR3 arrestin engagement and membrane localization compared to the antagonist. In addition to scavenging ligands to regulate canonical receptor function, ACKR3 outcompetes CXCR4 for arrestins^62^. This is thought to protect CXCR4 from downregulation and desensitization due to overstimulation by CXCL12^4,62,63^. By freeing arrestins basally engaged with ACKR3, while also blocking activation by CXCL12, the inverse agonists could act as tools to indirectly target CXCR4 or CXCR3 for downregulation.

All three nanobodies had profound inhibitory effects on the constitutive internalization of ACRK3. The atypical receptor undergoes both agonist-promoted internalization which is dependent on GRK phosphorylation as well as a ‘passive’ GRK-independent cycling between the plasma membrane and endosomes^5^. This second mechanism of receptor turnover is observed in other chemokine receptors^64^ and contributes to chemokine scavenging, but is insufficient to fully replace the active internalization response^4^. All three nanobodies retain the receptor at the plasma membrane, with the inverse agonists exhibiting significantly greater effects than the neutral antagonist in WT HEK293 cells. The nanobodies also retained ACKR3 at the plasma membrane in GRK2/3/5/6 KO cells, suggesting that even the neutral antagonist impacts the previously described passive internalization. These results clearly show that ACKR3 constitutive internalization occurs through both a GRK-dependent pathway, which requires receptor constitutive activation, and a GRK-independent pathway, operating via a heretofore undescribed mechanism. ACKR3 internalization does not require arrestins, which suggests a clathrin-independent mechanism, potentially through coordination via adenosine diphosphate ribosylation factors (ARFs), which regulate internalization and recycling pathways^65^. Alternatively, local membrane domains with specific lipid composition or curvature could sort GPCRs and mediate endocytosis during the natural turnover of the plasma membrane^66–68^. Thus, stabilization of ACKR3 by nanobody binding may segregate the receptor away from membrane regions primed to internalize, thereby leading to the trapping effect we observe.

The effects of VUN700 and VUN701 on basal cancer cell motility suggest that the constitutive activity of ACKR3 is implicated in migratory signaling. The degree of inhibition was greater for VUN700 than VUN701 in agreement with the efficacy of the inactivating responses to the two nanobodies. The cells tested also express CXCR4 and an attractive explanation for the inhibition might be that the cells secrete CXCL12 and the balance of chemokine scavenging by ACKR3 is needed for CXCR4-mediated migration. However, blocking CXCR4 with VUN400^39^ had no impact on the basal migration of these cells, indicating the motility observed is not due to CXCL12 stimulation of CXCR4. This therefore implies that inhibition of CXCL12 scavenging is not the mechanism for the impaired migration with ACKR3 nanobodies. We propose a chemokine-independent role for ACKR3 in basal motility of cancer cells. However, further studies are required to unravel how exactly and by which signaling pathways ACKR3 affects cell motility.

Biologics, including nanobodies, constitute an increasing proportion of FDA-approved therapeutics^41,69,70^ The inverse agonist nanobodies developed in this study may possess several features with therapeutic potential. Besides their antagonistic properties, the ability of these nanobodies to shift the receptor into an inactive conformation may also prevent crosstalk between ACKR3 and interacting proteins and receptors (like CXCR4, EGFR, Cx43^71–73^). As noted above, ACKR3 protects CXCR4 from desensitization^4^. Such a mechanism could have important implications for targeting this axis in cancer therapeutics. A dual-targeted approach could on one hand antagonize CXCR4 while also downregulate the canonical receptor via ACKR3 inverse agonism, as CXCL12 and arrestins would further desensitize CXCR4 and complement the direct inhibition. ACKR3 complexes with the gap junction protein Cx43 upon CXCL12 activation in astrocytes^73^. The atypical receptor coordinates the internalization of Cx43 in a β-arrestin-dependent manner, which inhibits gap intercellular communication. The inverse agonistic nanobodies could therefore preserve Cx43 on the plasma membrane and protect these structures. Knowing the capabilities of the current inverse agonists, expanding their modulatory activity through structural engineering would be interesting. Antibody engineering also generated a universal platform to support nanobody structural determination of membrane proteins^74^. New computational methods for designing nanobodies targeting specific epitopes with high affinity binders will continue to expand possible applications^75,76^. Further studies are necessary to ascertain whether differentially modulating ACKR3 is essential and/or beneficial when targeting ACKR3-related diseases including cardiovascular diseases (as atherosclerosis)^77^, autoimmune diseases (as multiple sclerosis)^78,79^, and cancer^57,80^.

In summary, we have identified a basal state of ACKR3 that displays constitutive activity that is involved in cancer cell motility. We investigated this through new nanobodies binding to the extracellular site of ACKR3 with unique properties. The inverse agonistic properties of two of these molecules emphasize the constitutive activity of the receptor by impairing basal β-arrestin engagement as well as its constitutive internalization by stabilizing an inactive receptor conformation. Inhibition of basal ACKR3 activity attenuated basal cell motility, which reveals a new role for ACKR3 in cell biology and cancer in particular. These results open new avenues and strategies for therapeutically targeting ACKR3.

## Methods

### Cell culture and transfection

The generation of the Clustered Regularly Interspaced Short Palindromic Repeats (CRISPR) genome-edited NanoLuc-ACKR3 Knock In (KI) HeLa cell line was described previously. Human embryonic kidney 293T (HEK293T) and MDA-MB-231 breast cancer cells were obtained from ATCC. HEK293 Parental and CRISPR HEK293 β-arrestin1/β-arrestin2 Knock Out (KO) cells were provided by Asuka Inoue from Tohoku University. Stable HEK293 expressing ACKR3 were kindly provided by Meritxel Canals from University of Nottingham. The stable, homogenous ACKR3 expression was necessary to resolve changes to the surface receptors by flow cytometry. HEK293 Parental and CRISPR GRK2/3/5/6 KO HEK293 cells were provided by Julia Drube and Carsten Hoffmann from University Hospital Jena. NanoLuc-ACKR3 HeLa, HEK293T, HEK293, HEK293 β-arrestin1/2 KO and stable ACKR3 HEK293 were cultured in Dulbecco’s Modified Eagle’s Medium (DMEM, Thermo Fisher Scientific, Gibco, #41966) supplemented with 100 Units of penicillin, 100 g/mL streptomycin (Pen/Strep, Gibco, #15140-122) and 10% (v/v) Fetal Bovine Serum (FBS, Bodinco). ACKR3 stable cell line was maintained under antibiotic selection with G418 500ug/ml (#A1720 Sigma-Aldrich).

HEK293T, HEK293 and HEK293 β-arrestin1/2 KO cells were transfected in suspension with a total of 1 μg DNA and 6 μg 25 kDa linear polyethyleimine (PEI, Polysciences Inc.) in 150 mM NaCl solution per 1 million cells. DNA encoding ACKR3 and biosensors was supplemented with empty pcDEF3 to obtain a total DNA amount of 1 μg. The DNA-PEI mixture was vortexed for 5 seconds and incubated for 15 min at room temperature (RT). Cells were detached with Trypsin (Gibco) and resuspended in DMEM. A 3×10^5^ cells/mL HEK293T cell suspension was added to DNA-PEI mixture and cells were seeded at a density of 30,000 in white flat-bottom 96-well plates (Greiner Bio-One) and incubated for 48h.

### ACKR3 Nanobodies selection via phage-display

Llama immunization, library construction, and nanobody selection were performed as described previously^81–83^. Briefly, two llamas were immunized with ACKR3-encoding plasmid DNA, cDNA was generated from peripheral blood mononuclear cells, nanobody sequences were amplified by PCR, and nanobody phage display libraries were constructed. Selections for ACKR3-specific binders were performed using three consecutive rounds of phage panning on ACKR3-expressing or empty (null) virus-like particles (Integral Molecular, Philadelphia, PA, USA) immobilized in MaxiSorp plates (Nunc, Roskilde, Denmark).

### Nanobodies production

Nanobody-FLAG-6×His proteins purification was performed as previously described^84^. pMEK222-transformed BL21 codon+ DH5alpha cells, respectively, were grown in LB/2% glucose O/N at 37 °C. The O/N preculture was inoculated in regular terrific broth and grown for 3 h at 37 °C, after which periplasmic expression of nanobodies was induced for another 4 h by addition of 1 mM isopropyl-b-D-thiogalactopyranoside (IPTG, Sigma-Aldrich) to the culture medium. After production, cultures were spun down for 30 min at 3500 × g and the pellets were frozen overnight at −20 °C. The next day, pellets were thawed and resuspended in PBS. The resuspended pellet was incubated at 4 °C for 2 h. Cultures were spun down for 30 min at 3500 × g at 4 °C and, after filtering using a 0.45 μM filter (VWR), the periplasmic fraction (supernatant) was stored at 4 °C until purification.

Nanobody-FLAG-6×His proteins were purified using ROTI®Garose-His/Co Beads (Carl Roth GmbH & Co, DE). Samples were first eluted in a buffer containing 500 mM imidazole PBS (Sigma-Aldrich, St. Louis, MO, USA) and after dialized using Snakeskin Dialysis Tubing 10 kDa molecular weight cut off (MWCO) membranes (Thermo Fisher Scientific) in phosphate-buffered saline (PBS). Purity of all produced and purified nanobodies was verified by sodium dodecyl sulfate-polyacrylamide gel electrophoresis (SDS-PAGE) under reducing conditions.

### NanoBRET CXCL12 competition assay

30k cells per well of NanoLuc-ACKR3 Knock In (KI) HeLa cells were seeded in white flat-bottom 96-well plate. 24 h later, cells were washed with PBS once, and Hank’s Buffered Saline Solution (HBSS) supplemented with 0.1% bovine serum albumin (BSA), 10 mM HEPES, 1 mM MgCl_2_ and 2 mM CaCl_2_ was added to the cells. Subsequently, increasing concentrations of unlabelled CXCL12 (Almac) or unlabelled nanobodies were added as indicated in the figures, and incubated for 45 min at RT. Next, 3.3 nM CXCL12-AF647 (Almac) was added and incubated for 15 min at RT. Next, luciferase substrate (Furimazine, Nano-Glo® substrate (Promega, #N1110, final concentration of 15 μM)) was added and luminescence was measured using a PheraSTAR plate reader (BMG) with 460 ± 80 nm and 610-LP nm filters. BRET data were normalized to full homologous displacement (0%) and fluorescent CXCL12 only (100%). Data were analyzed with a nonlinear fit to create a dose-response curve in Graph Pad Prism Version 10.2.0. Data from all independent experiments were used in the analysis and calculation of standard deviation.

### NanoBRET assays

1 million cells were transfected with 2 μg total DNA, consisting of BRET donor, BRET acceptor, supplemented with empty plasmid. Position of genetically fused sensors is given when stating the construct (e.g. in Nluc-ACKR3 Nluc tag is located in the N-terminus of ACKR3, while ACKR3-Nluc, Nluc is on the C-terminus). In the β-arr1/2 recruitment and internalization experiments, 30-50 ng of HA-ACKR3 WT-Nluc was used in combination with 150-250ng of the different acceptors, β-arr1/2-mVenus, mVenus-CAAX or Rab5a-mVenus (donor:acceptor ratio 1:5). Cells were transfected in suspension with PEI in a ratio of PEI ug:DNA ug 6:1, with a total amount of 2ug of DNA per million cells. Cell suspension of 300k/ml is used to seed 30k/well cells in 96 well plate. 48h after transfection, cells were washed with PBS once, and HBSS supplemented with 0.1% BSA, 10 mM HEPES, 1 mM MgCl2 and 2 mM CaCl2 was added to the cells. Incubation with the luciferase substrate (Furimazine, Nano-Glo® substrate (Promega, #N1110, final concentration of 15 μM)) followed and the basal BRET was measured for 5 minutes. BRET for this pair Nluc and mVenus was measure at 460 ± 30 nm and 535 ± 30 nm respectively. Next, cells were treated with CXCL12 or nanobodies indicated in the legend of the figures and BRET was measured for 60 minutes. Normalized BRET ratio was then calculated by dividing the raw BRET values of each well from the ligand induced results by the basal BRET measured before stimuli (baseline). For vehicle normalization, normalized BRET value was divided by the vehicle condition (without ligand) over time, to normalize for effects of the drop in furimazine availability. Data were analyzed with a nonlinear fit (three parameters model, with equation 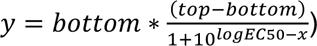 to create)*)+ a dose-response curve in GraphPad Prism. Data from all independent experiments were used in the analysis and calculation of standard deviation. Graph Pad Prism Version 10.2.0.

### Intramolecular FlAsH-NanoBRET assays

β-arrestin2 conformational change biosensors used in this work were previously described^52^. CRISPR control cells were transfected with 1.2 μg of untagged ACKR3, 0.12 μg of a β-arrestin2 FlAsH-tagged biosensor C-terminally coupled to NanoLuc, and 0.25 µg of an empty vector, following the Effectene transfection reagent protocol by Qiagen. In total, three sensors were used, numbered as β-arrestin2 FlAsH-3,5 10-NanoLuc sensors. The following day, 40,000 cells were seeded per well into poly-D-lysine-coated 96-well plates and incubated overnight at 37 °C. For this study, the FlAsH (fluorescein arsenical hairpin-binder)-labeling procedure, previously described by Hoffmann et al.^85^, was adjusted for 96-well plates. Briefly, the cells were washed twice with PBS, then incubated with 250 nM FlAsH or mock DMSO in labeling buffer (150 mM NaCl, 10 mM HEPES, 25 mM KCl, 4 mM CaCl_2_, 2 mM MgCl_2_, 10 mM glucose; pH 7.3), complemented with 12.5 μM 1,2-ethane dithiol (EDT) for 1 hour at 37 °C. After aspiration of the FlAsH labeling and mock labeling solutions, the cells were incubated for 10 min at 37 °C with 100 μl of 250 μM EDT in labeling buffer, per well. The NanoLuc substrate was added and a basal measurement was recorded for 3 min. Subsequently, either CXCL12 or VUN700, VUN701 or VUN702 nanobodies were added in the required concentrations and BRET was measured for 20 minutes. Analysis of the BRET change was performed as described above (see Section “NanoBRET assays”). Measurements were performed using the Synergy Neo2-provided BRET2 filter (Emission wavelengths 400/510).

### Expression and purification of ACKR3 (HDX)

For production in insect cells, the full-length gene of human ACKR3 was subcloned into pFastBac1 to enable infection of sf9 insect cells. The construct bore a hemagglutinin signal peptide followed by a Flag-tag preceding the receptor sequence. ACKR3 N13, N22 and N33 residues were substituted with a Glutamine in order to avoid N-glycosylation.

Flag-ACKR3 was expressed in sf9 insect cells using the pFastBac baculovirus system (Thermo Fisher Scientific). Cells were grown in suspension in EX-CELL 420 medium (Sigma-Aldrich) and infected at a density of 4 x 10^6^ cells/ml with the recombinant baculovirus. Flasks were shaken for 48 h at 28 °C, subsequently harvested by centrifugation (3,000 x g, 20 min) and stored at -80 °C until usage. Cell pellets were thawed and lysed by osmotic shock in a buffer containing 10 mM Tris (pH 7.5), 1 mM EDTA, 2 mg/ml iodoacetamide, 1 μM ACKR3 agonist VUF11207 and protease inhibitors: 50 μg/ml Leupeptin (Euromedex), 0.1 mg/ml Bensamidine (Sigma-Aldrich), and 0.1 mg/ml Phenylmethylsulfonyl fluoride (PMSF; Euromedex). Lysed cells were centrifuged for (38,400 x g, 10 mins) and the resulting pellet was solubilised and dounce-homogenised 20 x in buffer containing 50 mM Tris (pH 7.5), 150 mM NaCl, 2 mg/ml iodoacetamide, 1 μM VUF11207, 0.5% (w/v) dodecyl maltoside (DDMm Anatrace), 0.1% (w/v) cholesteryl hemisuccinate (CHS) and protease inhibitors. The homogenate was subsequently stirred for 1 h at 4 °C and centrifuged (38,400 x g, 30 min). The supernatant was then loaded onto M2 anti-Flag affinity resin (Sigma-Aldrich) using gravity flow. Resin was subsequently washed with 10 column volumes (CV) of DDM wash buffer containing 50 mM Tris, 150 mM NaCl, 0.1 μM VUF11207, 0.1% (w/v) DDM, 0.02% (w/v) CHS. Detergent was then gradually exchanged from DDM to lauryl maltose neopentyl glycol (MNG, Anatrace) using decreasing ratios of DDM wash buffer and buffer containing 50 mM Tris, 150 mM NaCl, 0.02 μM VUF11207, 0.2% (w/v) MNG, 0.05% (W/V) CHS. Once detergent was fully exchanged, MNG and CHS concentration were steadily reduced to 0.005% and 0.001% respectively. ACKR3 was finally eluted in 50 mM Tris, 150 mM NaCl, 0.02 μM VUF11207, 0.005% (w/v) MNG, 0.001% (w/v) CHS and 0.4 mg/ml Flag peptide (Covalab). The eluate was concentrated using a 50 kDa MWCO concentrator (Millipore), then ACKR3 was purified by size exclusion chromatography (SEC) using a Superdex 200 Increase (10/300 GL column) connected to an ÄKTA purifier system (GE Healthcare) and eluted in buffer elution buffer without Flag peptide or VUF11207. Fractions containing monomeric ACKR3 were concentrated to between 20 and 25 μM, aliquoted, flash-frozen and stored at -80 °C prior to HDX experiments.

### HDX-MS experiments

HDX-MS experiments were performed using a Synapt G2-Si HDMS coupled to nanoAQUITY UPLC with HDX Automation technology (Waters Corporation). ACKR3 in LMNG detergent was concentrated up to 20-25 µM and optimization of the sequence coverage was performed on undeuterated controls. Various quench times and conditions were tested; in the presence or absence of different denaturing or reducing reagents with or without longer trapping times to wash them out. The best sequence coverage and redundancy for ACKR3 were systematically obtained without the addition of any denaturing agents in the quench buffer. Mixtures of receptor and nanobody were pre-incubated to reach equilibrium prior to HDX-MS analysis. Analysis of freshly prepared ACKR3 apo, ACKR3: nanobody (1: 2 ratio) were performed as follows: 3 µL of sample are diluted in 57 µL of undeuterated for the reference or deuterated last wash SEC buffer. The final percentage of deuterium in the deuterated buffer was 95%. Deuteration was performed at 20 °C for 0.5, 2, 5, 30 and 120 mins. Next, 50 µL of reaction sample are quenched in 50 µL of quench buffer (50 mM KH_2_PO_4_, 50 mM K_2_HPO_4_, 200 mM tris(2-carboxyethyl)phosphine (TCEP) pH 2.3) at 0 °C. 80 µL of quenched sample are loaded onto a 50 µL loop and injected on a Nepenthesin-2 column (Affipro) maintained at 15 °C, with 0.2% formic acid at a flowrate of 100 µL/min. The peptides are then trapped at 0 °C on a Vanguard column (ACQUITY UPLC BEH C18 VanGuard Pre-column, 130Å, 1.7 µm, 2.1 mm X 5 mm, Waters) for 3 min, before being loaded at 40 µL/min onto an Acquity UPLC column (ACQUITY UPLC BEH C18 Column, 1.7 µm, 1 mm X 100 mm, Waters) kept at 0 °C. Peptides are subsequently eluted with a linear gradient (0.2% formic acid in acetonitrile solvent at 5% up to 35% during the first 6 min, then up to 40% and 95% over 1 min each) and ionized directly by electrospray on a Synapt G2-Si mass spectrometer (Waters). Maldi Imaging High Definition MSE (HDMSE) data were obtained by 20-30 V trap collision energy ramp. Lock mass accuracy correction was made using a mixture of leucine enkephalin and GFP. For every tested condition we analyzed two to three biological replicates, and deuteration timepoints were performed in triplicates for each condition.

Peptide identification was performed from undeuterated data using ProteinLynx global Server (PLGS, version 3.0.3, Waters). Peptides are filtered by DynamX (version 3.0, Waters) using the following parameters: minimum intensity of 1000, minimum product per amino acid of 0.2, maximum error for threshold of 10 ppm. All peptides were manually checked, and data was curated using DynamX. Back exchange was not corrected since we are measuring differential HDX and not absolute one. Statistical analysis of all ΔHDX data was performed using Deuteros 2.048 and only peptides with a 99% confidence interval were considered.

### pFastBac constructs and mutant generation (NMR)

Human ACKR3 WT pFastBac1 plasmid was generously provided by Dr. Tracy Handel (UC San Diego). The construct is comprised of a gp64 promoter, N-terminal HA signal sequence for membrane localization, human WT ACKR3 (unmodified expect with removal of the N-terminal methionine), a C-terminal PreScission protease cleavage tag, and FLAG / 10× His tags as described previously^86^.

### Baculovirus preparation and ACKR3 expression (NMR)

Baculovirus generation and ACKR3 expression was performed as described previously^86,87^. Briefly, recombinant baculovirus was produced using the Bac-to-Bac Baculovirus Expression System (Invitrogen). A pFastBac1 plasmid containing the described ACKR3 construct was transformed into DH10Bac *E. coli* (Thermo Fisher) and subsequently plated onto LB agar with 50 μg/ml, kanamycin, 7 μg/ml gentamicin, 10 μg/ml tetracycline, 100 μg/ml Bluogal and 40 μg/ml, IPTG (Teknova). Blue/white screening identified recombinant (white) colonies, of which individual clones were inoculated in 5mL of LB with 50 μg/ml ^-1^, kanamycin, 7 μg/ml ^-1^ gentamicin, and 10 μg/ml ^-1^ tetracycline and placed in a 37 °C shaking incubator overnight. Cultures were pelleted, lysed, and neutralized using buffers from the GeneJET Plasmid Midiprep Kit (Thermo Fisher) and bacmid was purified by isopropanol precipitation (see reference^86^ for details). Final bacmid pellets were solubilized in 40 mM Tris, pH 8 and 1 mM EDTA. To transfect Sf9 cells a mixture of purified bacmid (5 μl – 1 µg total DNA), X-TremeGENE HP DNA (3 μl) and Expression Systems Transfection Medium (100 μl) was mixed and added to 2.5 ml of Sf9 cells at ∼ 1.2 x 10^6^ cells/ml and the bacmid-cell mixture was placed in a 24-well, deep-well plate (Thomson Instrument Company) covered with a polyurethane sealing film (Diversified Biotech). Cells were incubated at 27 °C for 96 h at 300 rpm. Cells were subsequently pelleted, and the supernatant was collected to isolate P0 (“zero passage”) virus. P0 virus titers were determined using gp64 titer assay as described previously^86^. Next, P1 (“first passage”) virus was produced by infecting 50 ml of Sf9 cells at a density of ∼ 2.0 x 10^6^ cells/ml with titered P0 virus at an MOI of 0.1-0.5. Cells were incubated at 27 °C for 72 h shaking at 144rpm. Cells were pelleted and the supernatant was collected and titered using the same gp64 titering assay^86^. Large scale expression of labeled ACKR3 was performed by adding high titer P1 virus (≥ 1 x 10^-9^ IU/ml) to ∼2 L of Sf9 cells in methionine deficient medium (Expression Systems) at a density of 3.5-4.0 x 10^6^ cells/ml at an MOI of 5^87^. After 5 hours post-infection, 250 mg/L ^13^CH_3_-methionine (Cambridge Isotope Laboratories) was added to cells. Cells were incubated at 27 °C for 48 h then pelleted and stored at -80 °C.

### ACKR3 purification (NMR)

Receptor was purified described previously^87^ with some modifications. Briefly, frozen cell pellets (∼50 ml frozen cells per 2 L of cell culture) were diluted 1:1 in hypotonic buffer (10 mM HEPES pH 7.5, 20 mM KCl, 10 mM MgCl_2_, Roche Complete Protease Inhibitor Cocktail, 2 mg/ml iodoacetamide) and thawed on ice. Direct solubilization was performed by passing the cell slurry through a 16-gauge needle four times to aid solubilization by lysing cells. Cell slurry was added to solubilization buffer (100 mM HEPES pH 7.5, 800 mM NaCl, 1.5% (w/v) MNG, 0.3% (w/v) CHS) and incubated at 4 °C for 4 h with stirring. The mixture was spun down at 50,000 x g for 30 min and the supernatant (∼200 ml) was transferred to 4 x 50 ml conical tubes. 4 ml TALON cobalt resin slurry (Takara Bio Inc.) was added (1 ml/tube) with 10mM imidazole final concentration to limit non-specific binding, and the supernatant mixture was rocked overnight at 4 °C. This mixture was added to columns the following day, and cobalt resin was washed with 20 ml of two wash buffers (Wash Buffer 1: 50 mM HEPES pH 7.5, 400 mM NaCl, 0.1% (w/v) MNG, 0.02% (w/v) CHS, 10% glycerol, 20 mM imidazole; Wash Buffer 2: 50 mM HEPES pH 7.5, 400mM NaCl, 0.025% (w/v) MNG, 0.005% (w/v) CHS, 10% glycerol, 10 mM imidazole). ACKR3 was eluted with a high imidazole buffer (50 mM HEPES pH 7.5, 400 mM NaCl, 0.025% (w/v) MNG, 0.005% (w/v) CHS, 10% glycerol, 250 mM imidazole. Elutions were concentrated to 500 μl using a 30,000 MWCO Amicon Ultra-4 Centrifugal Filter Unit (Millipore Sigma) and buffer exchanged into Exchange Buffer (25 mM HEPES pH 7.5, 150 mM NaCl, and 0.025% (w/v) MNG, 0.005% (w/v) CHS) using a PD-10 desalting column (GE). Precission Protease and PNGaseF were added to purified ACKR3 overnight. The next day, 500 μl TALON cobalt resin was added, and the mixture was incubated with rocking for 2 h at 4 °C. The mixture was added to a new column to separate cleaved receptor from the tag-bound cobalt resin and washed with Exchange Buffer to collect the flow through. Flow through was concentrated to ∼1 ml before quantifying.

### Nuclear magnetic resonance (NMR)

Purified protein samples concentrated to ∼350 μl with 10% D_2_O by volume were loaded into a 5 mm Shigemi microtube. Heteronuclear single quantum coherence (HSQC) spectra were collected on a Bruker Avance 800 MHz spectrometer equipped with a triple-resonance cryogenic probe with experiments collected at 310 K. Experimental times were 27 h. Data were processed using NMRPipe^88^ and visualized in XEASY^89^. CXCL12 used in this assay was purchased from Protein Foundry, L.L.C.

### Expression and Purification of VUN700, VUN701, VUN702 (NMR)

The sequences of VUN700, VUN701, VUN702 were codon-optimized for *E. coli* expression and ordered from GenScript. The nanobodies were cloned into a pET28a-6×His-SUMO3 vector and expressed in BL21 DE3 *E.coli*. Cells were expressed at 37 degrees C in Luria-Bertani (LB) medium and induced with 1 mM IPTG at an OD600 of 0.8. Cultures continued to grow for 5 and a half hours before bacteria were pelleted by centrifugation and stored at -20°C. Bacterial pellets were resuspended in ∼20 mL of Buffer A (50 mM Na_2_PO_4_ (pH 8.0), 300 mM NaCl, 10 mM imidazole, 1 mM PMSF, and 0.1% (v/v) 2-mercaptoethanol (BME)) per pellet and lysed via sonication. Lysed cells were clarified at 18,000 x g and the supernatant was discarded. Pellets were resuspended by sonication in ∼20 mL of Buffer AD (6 M guanidinium, 50 mM Na_2_PO_4_ (pH 8.0), 300 mM NaCl, 10 mM imidazole) and spun down at 18,000 x g for 20 min. Using an AKTA-Start system (GE Healthcare), the supernatant was loaded onto a Ni-NTA column equilibrated in Buffer AD. The column was washed with Buffer AD, and proteins were eluted using Buffer BD (6 M guanidinium, 50 mM sodium acetate (pH 4.5), 300 mM NaCl, and 10 mM imidazole). Proteins were refolded overnight via drop-wise dilution into a 10-fold greater volume of Refold Buffer (50 mM Tris (pH 7.6), 150 mM NaCl) with the addition of 30 mM cysteine, and 1 mM cystine. Refolded protein was concentrated in an Amicon Stirred Cell concentrator (Millipore Sigma) using a 10 kDa membrane. Concentrated protein was added to 6-8 kDa dialysis tubing with the addition of ULP1 to cleave the N-terminal 6×His-SUMO3-tag and dialyzed at 25 °C against Refold Buffer overnight. The AKTA-Start system was used to load the cleaved protein onto a Ni-NTA column equilibrated in VUN701 Buffer A (Refold Buffer + 10 mM Imidazole). The column was washed with VUN701 Buffer A, and the protein was eluted using VUN701 Buffer B (Refold Buffer + 500 mM Imidazole). VUN701 underwent four rounds of dialysis in 5 mM ammonium bicarbonate, lyophilized, and stored at -80°C for further use. The purity and identity of nanobodies were confirmed by electrospray ionization mass spectrometry using a Thermo LTQ instrument and SDS-PAGE with Coomassie staining.

### Flow cytometry

HEK293 cells stably expressing ACKR3 were washed with PBS and lifted with Accumax (Invitrogen). 150k cells were transferred per well of conical 96 well plates (Greiner). Cells were washed with cold FACS buffer (0.5% (w/v) BSA in PBS) and treated with corresponding ligands (316 nM of VUN700, VUN701 or VUN702, or 100 nM CXCL12)or vehicle (untreated) over 60 min at 37 °C with Assay media (0.5% (w/v) BSA, 25 mM HEPES in DMEM). After treatments, cells were treated with ice-cold buffers and kept on ice until readout. Cells were washed once with FACS buffer, followed by 2 acidic washes (0.2 M acetic acid, 0.5 M NaCl). Cells were washed 3 times with FACS buffer and labeled with 10 µL/10^6^ cells of PE-conjugated anti-ACKR3 antibody (11G8-PE, #FAB4227 R&D Systems) in FACS buffer for 1 h at 4 °C. Unbound antibody was then 3× washed away with FACS buffer. Surface ACKR3 was assessed by flow cytometry using a Guava easyCyte benchtop flow cytometer (Luminex). The mean fluorescence intensity (MFI) representing the amount of surface labeling for each experiment was quantified using Floreada Software (Floreada.io). The constitutive internalization was then represented by the ratio of the MFI of the treated samples to the non-treated controls, over each time point. Graph Pad Prism Version 10.2.0 (335), multiple comparisons two-way ANOVA Tukey test (*p<0.05).

### Basal motility

MDA-MB-231 metastatic breast cancer cells were detached from a subconfluent flask and plated at high density in the central imaging chambers of ‘Ibidi µ-Slide Chemotaxis’ and allowed to attach. The central imaging chamber was flushed twice with (-/+) 100 nM nanobody-containing media, and then reservoirs filled with the same treatment, giving a stable uniform concentration over the cells throughout the experiment. 1 h after the commencement of treatment, cells were imaged on the Nikon Ti2 microscope at 10X objective within a controlled chamber of 37 °C and 5% CO_2_, using a motorized stage to image each central chamber every 30 min for 16 h. The migration of 40 randomly chosen cells in each group were manually tracked using the ImageJ/Fiji Plugin ‘Manual Tracking’, and these tracks were compiled and analyzed for their trajectories by the plugin ‘Chemotaxis and migration tool’ (Ibidi). Accumulated distance (total distance travelled), Euclidean distance (straight line distance from cell starting point to end point) and velocity (accumulated distance/time) were analyzed. The data shows the mean +/-SD of each group relative to their internal control, combined from multiple independent repeats. Significance determined by one-way ANOVA Dunnett test (p<0.05(*), 0.005 (**), 0.0005 (***), <0.0001 (****)).

### Data availability

The HDX mass spectrometry data have been deposited to the ProteomeXchange Consortium via the PRIDE partner repository with the dataset identifiers PXD051149.

## Acknowledgements

Research was supported by the ONCORNET 2.0 (ONCOgenic Receptor Network of Excellence and Training 2.0) PhD training programme funded by the European Commission for a Marie Sklodowska Curie Actions (H2020-MSCA grant agreement 860229) and by a National Institutes of Health Allergy and Infectious Disease grant (NIAID R37AI058072 to BFV). HDX-MS experiments were carried out using the facilities of the Montpellier Proteomics Platform (PPM, BioCampus Montpellier). CC was supported by the ZonMw Veni grant 09150162010212. CB acknowledges funding from the regional funds FEDER/Région Occitanie, MUSE, Labex EpiGenMed and the French Agence Nationale de la Recherche (project LEUKOCEPTOR ANR-21-CE44-0007-01). CTS and MJS acknowledge funding from the European Innovation Council through its Horizon Europe Pathfinder Open programme (Grant Agreement No. 101131014) and the Swiss State Secretariat for Education, Research and Innovation (SERI).

