## Supplementary information for "Constitutive activity of an atypical chemokine receptor revealed by inverse agonistic nanobodies"

**Supplementary Table 1.** pEC50s and pIC50s (± SD) of CXCL12 and ACKR3-targeting Nbs for BRET experiments.

| | **CXCL12** | **VUN700** | **VUN701** | **VUN702** |
|---|---|---|---|---|
| **CXCL12 displacement** | 9.9± 0.1 | 8.5 ± 0.1 | 8.7 ± 0.1 | 8.1 ± 0.2 |
| **β-arrestin1 recruitment** | 8.7 ± 0.2 | 8.2 ± 0.3 | n.a. | 8.1 ± 0.2 |
| **β-arrestin2 recruitment** | 8.9 ± 0.1 | 8.0 ± 0.2 | n.a. | 7.8 ± 0.1 |
| **Internalization CAAX** | 9.1 ± 0.1 | 8.1 ± 0.1 | 8.3 ± 0.2 | 7.5 ± 0.1 |
| **Early endosomes Rab5a** | 10.0 ± 0.2 | 7.0 ± 0.2 | n.a. | 7.2 ± 0.3 |
| **Parental Intern. CAAX** | 9.7 ± 0.2 | 8.3 ± 0.1 | 8.4 ± 0.1 | 8.1 ± 0.1 |
| **dQ KO Intern. CAAX** | n.a. | 8.2 ± 0.1 | 8.4 ± 0.1 | 8.0 ± 0.0 |
| **Parental β-arrestin2 recruitment** | 9.0 ± 0.3 | n.a.* | n.a. | n.a.* |
| **dQ KO β-arrestin2 recruitment** | n.a. | n.a. | n.a. | n.a. |
| **Parental Intern. CAAX** | 8.9 ± 0.1 | 7.8 ± 0.1 | 8.0 ± 0.2 | 7.4 ± 0.1 |
| **β -arr2 KO Intern. CAAX** | 9.0 ± 0.1 | 7.8 ± 0.1 | 8.2 ± 0.2 | 7.6 ± 0.1 |

n.a. (not applicable), *window too small

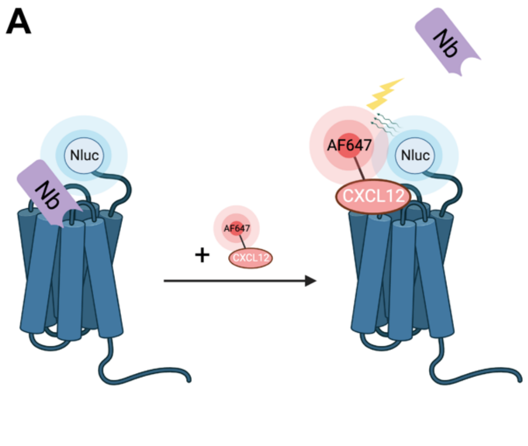


**Supplementary Figure 1. All ACKR3-targeting nanobodies displace CXCL12 with comparable affinities. A)** Schematic illustration of NanoBRET-based competition binding assay between ACKR3-targeting nanobodies and CXCL12-AF647. **B)** Displacement curves of unlabeled CXCL12 (blue circle), or nanobodies VUN700 (green triangle), VUN701 (purple inverted triangle) or VUN702 (yellow diamond) by CXCL12-AF647 (3.3 nM) binding to Nanoluc-ACKR3 CRISPR Knock In HeLa cells. Data are shown as the average of three independent experiments performed in triplicate ± SD, normalized to maximum CXCL12-AF647 binding.

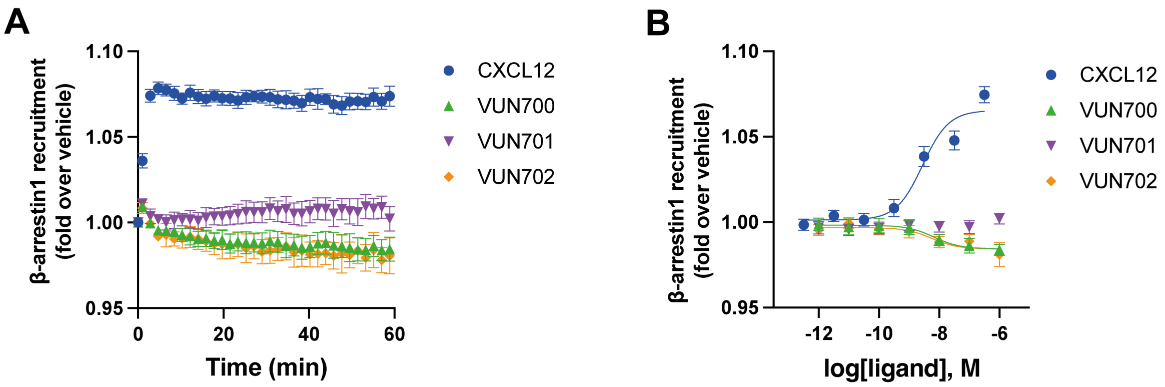


**Supplementary Figure 2. ACKR3 nanobodies acts as inverse agonists and neutral modulators for β-arrestin1 recruitment. A-B)** Recruitment of β-arrestin1-mVenus to ACKR3-Nluc measured by BRET. **(A)** The time-dependent change in BRET over 60 min after treatment with either 316 nM of CXCL12 (blue circle) or 1 μM of VUN700 (green triangle), VUN701 (purple inverted triangle), or VUN702 (yellow diamond). **(B)** Dose response curves of nanobodies or CXCL12 at 60 min, at 37 °C in HEK293T cells. Data is shown as the average ± SD of three independent experiments performed in duplicates.

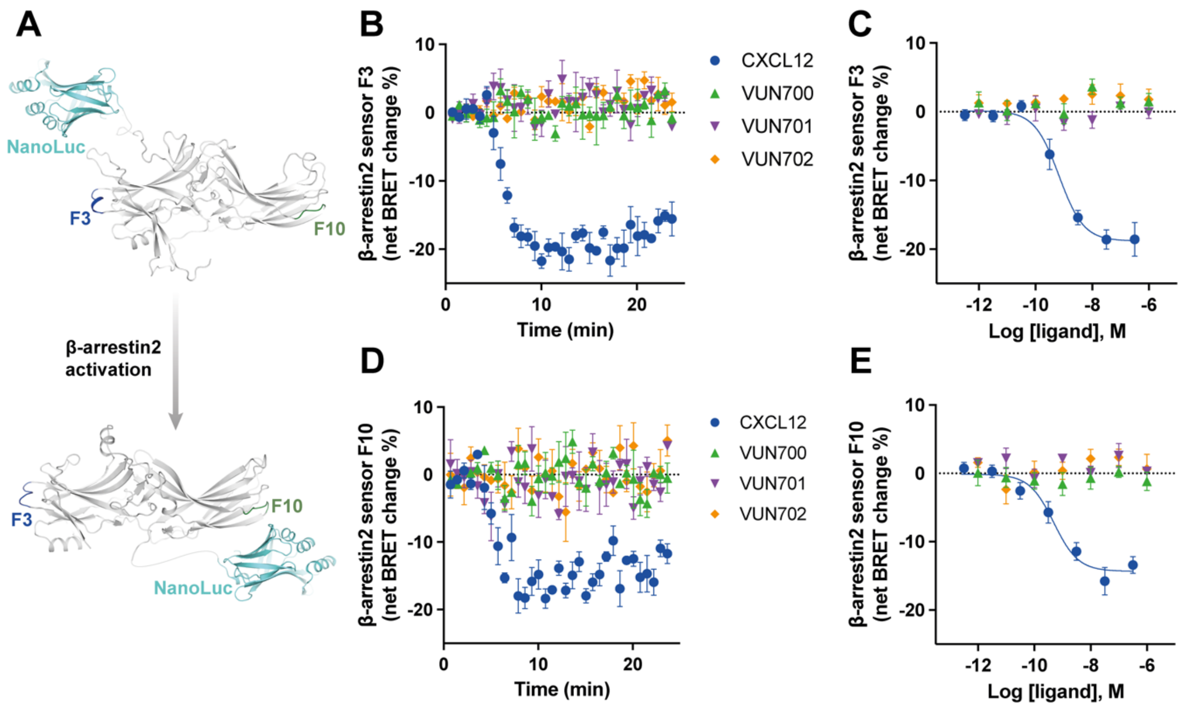


**Supplementary Figure 3. Nanobody treatment of ACKR3 does not lead to a change in β-arrestin2 conformational biosensors F3 and F10 BRET. A)** Schematic illustration of BRET-based FlAsH-tagged (CCPGCC) sensors F3 (between 48 and 49) and F10 (between 262 and 263) on β-arrestin2. **B-C)** F3 β-arrestin2 NanoBRET conformational biosensor with ACKR3 **(B)** time-dependent change in BRET over 25 min with either 316 nM of CXCL12 (blue circle) or 1 µM of VUN700 (green triangle), VUN701 (purple inverted triangle) or VUN702 (yellow diamond) and **(C)** DRCs effect of nanobodies at 25 min, at 37 °C in HEK293 cells. **D-E)** F10 β-arrestin2 NanoBRET conformational biosensor with ACKR3 **(D)** kinetics over 25 min with either 316 nM of CXCL12 or 1 µM of VUN700, VUN701 or VUN702 and **(E)** DRCs effect of nanobodies at 25 min, at 37 °C in HEK293 cells. Data is shown as the average ± SD of three independent experiments performed in triplicates.

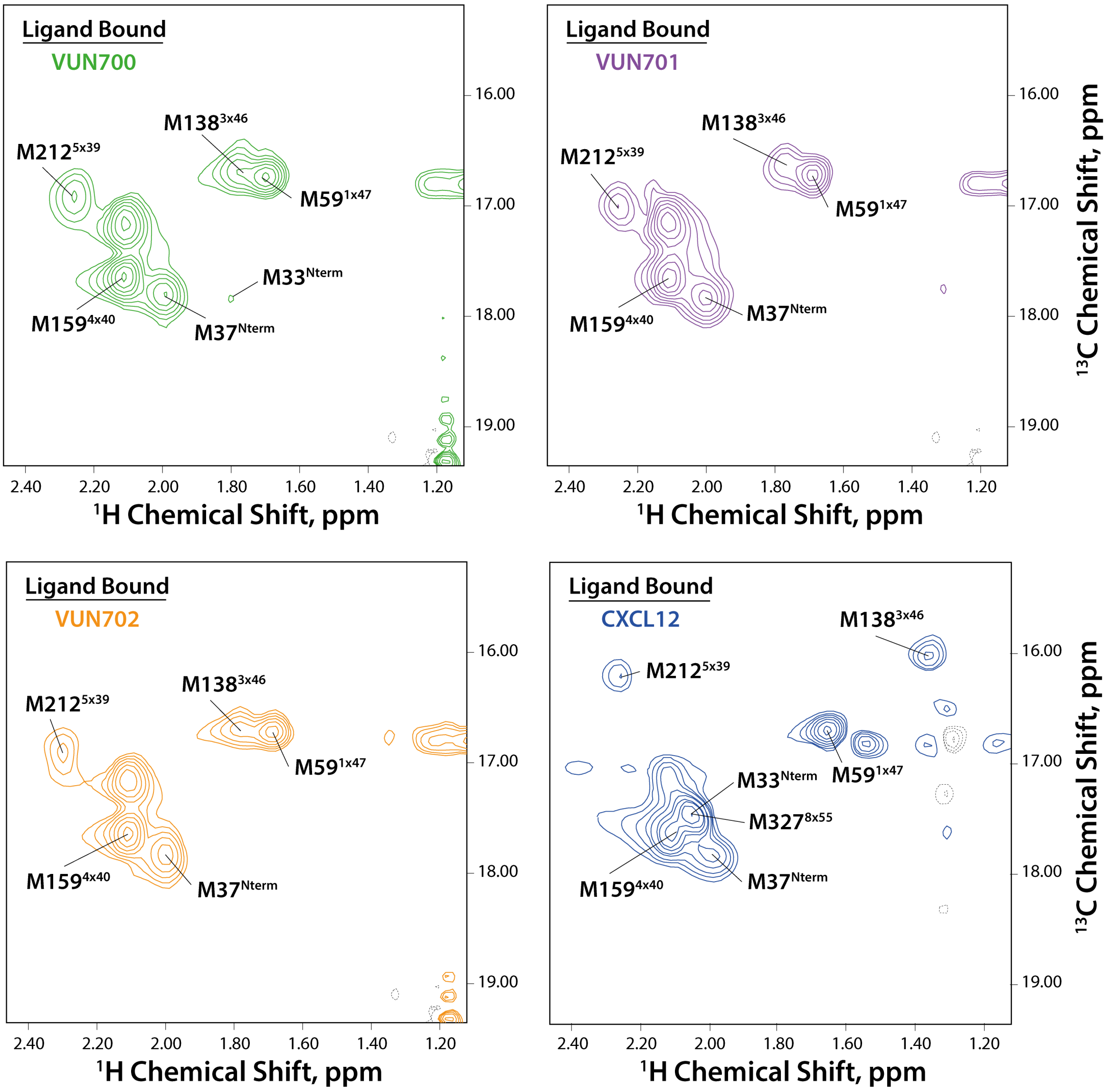


**Supplementary Figure 4. NMR spectra of ACKR3 with nanobodies VUN700, VUN701, VUN702, and CXCL12.** $^{1}$H-$^{13}$C heteronuclear single quantum coherence NMR spectra of WT-ACKR3 with different ligands (three ACKR3-binding nanobodies and CXCL12[1]) at 310 K.

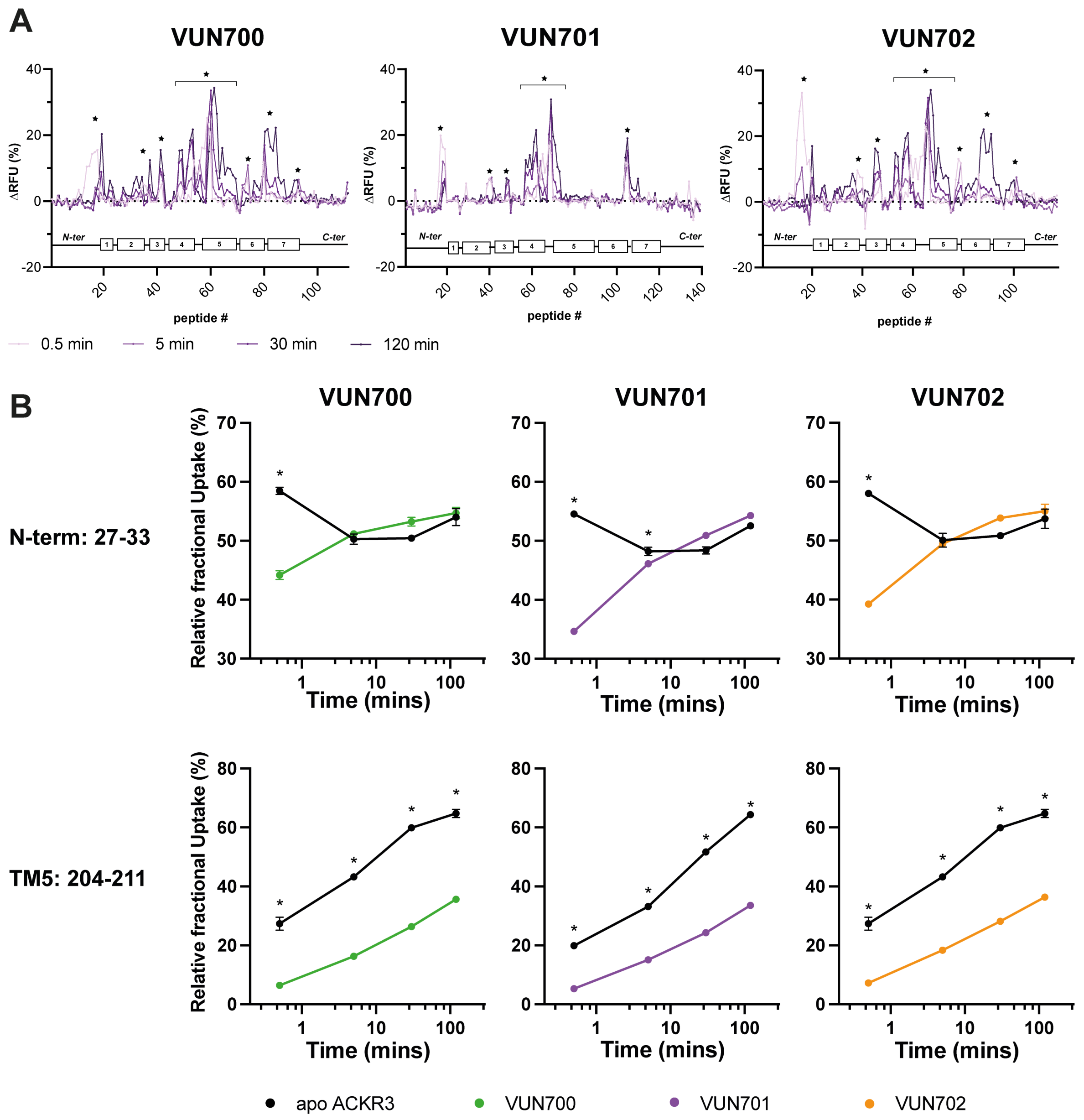


**Supplementary Figure 5. HDX data of ACKR3 with nanobodies VUN700, VUN701, and VUN702. A)** Butterfly plots showing time-dependent change in ΔRFU for identified ACKR3 peptides in both apo and bound forms. **B)** HDX uptake plots showing time-dependent change in relative fractional uptake for ACKR3 peptides at the N-terminal and TM5 levels upon nanobodies binding. Uptake represents the average and SD of three technical replicates from one biological preparation of ACKR3. Data is representative of three biological replicates. Statistically significant changes were determined using Deuteros 2.0 software[2] (*, $p \leq 0.01$).

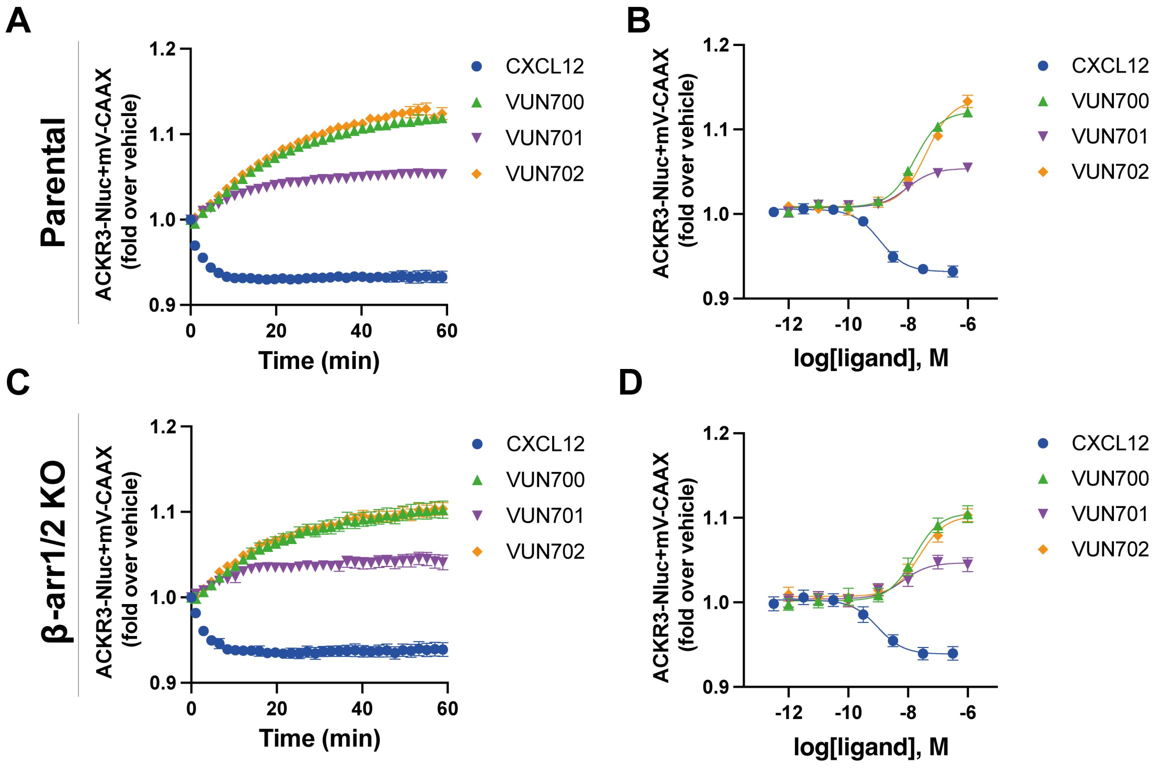


**Supplementary Figure 6. ACKR3 nanobodies retain ACKR3 at the plasma membrane by a β-arrestin1/2 independent mechanism. A-D)** Internalization measured by BRET in agonist mode, between donor ACKR3-Nluc and mV-CAAX in **(A, B)** HEK293 or in **(C, D)** HEK293 β-arrestin 1/2 CRISPR KO cells. **(A, C)** Time-dependent change in BRET over 60 min with either 316 nM of CXCL12 (blue circle) or 1 µM of VUN700 (green triangle), VUN701 (purple inverted triangle), or VUN702 (yellow diamond) and **(B, D)** dose response curves of CXCL12 or nanobodies at 60 min, at 37°C in HEK293T cells. Data is shown as the average ± SD of four independent experiments performed in triplicates.
